# Ecological and dietary differences between Ugandan chimpanzee communities with possible implications on tool use

**DOI:** 10.1101/670539

**Authors:** Iris Berger, Catherine Hobaiter, Matthew Bell, Delphine De Moor, Thibaud Gruber

## Abstract

Some East African chimpanzee (*Pan troglodytes schweinfurthii*) communities, such as the Sonso chimpanzees, display an unusually limited range of tool-use, but it remains unclear whether this is due to ecological and/or cultural factors. Information on ecological conditions and the diet of the Sonso chimpanzees in relation to neighbouring communities is needed. Here, we studied three adjacent communities in Budongo Forest (Sonso, Waibira, and Kamira), and the presumed core area of an undescribed community (Mwera), in the neighbouring Bugoma Forest. Through line-transects, we investigated (i) whether there were differences in food diversity and abundance between the communities’ home ranges; (ii) whether the home ranges differed in abundance of sticks and insect nests; and (iii) whether Sonso and Mwera chimpanzees differed in their diet (using faecal samples). Across communities, Sonso had the richest food availability and the lowest insect nest abundance. However, food availability in Mwera, Bugoma, was richer than Budongo communities that neighbour the Sonso territory, suggesting that there may be variation within Budongo. Data from faecal samples replicated our direct observations of food availability suggesting that Sonso chimpanzees had a broader diet than Mwera chimpanzees. This difference in foods availability may partially explain the Sonso chimpanzees’ lack of stick-tool-use, and low levels of insectivory. The tool repertoire of the other communities is currently unknown; however, we make predictions based on our ecological data. More detailed knowledge of small-scale variation in ecology within and between forest habitats may be important to advancing our understanding of the drivers of tool-use.

**SIGNIFICANCE STATEMENT – HIGHLIGHTED STUDENT PAPER:** To advance our knowledge of the role of ecological factors in the emergence of tool use in chimpanzees, a nuanced understanding of the ecological conditions different chimpanzee communities experience is needed. We studied four Ugandan chimpanzee communities in two forests. One of these communities, Sonso, in the Budongo Forest, is well-known for its restricted range of tool types, including a total absence of stick use. Food diversity and abundance were highest, and stick tool use opportunities (abundance of sticks and insect nests) were lowest for the core-habitat of the Sonso chimpanzees in contrast to the other communities. We argue that ecological factors play a role in their unusual pattern of tool use, and make predictions about the expected types of tool use in the other communities based on their ecology. Thus, our study provides information that may help advance our understanding of how tool use arises under varied socioecological circumstances.

## INTRODUCTION

In a landmark study, Whiten et al. (1999) showed that 39 behavioural patterns, mainly in the domain of tool use, were customary in some communities, but not in others. While the original article did not discuss ecological explanations in detail, much debate has followed regarding the impact of ecological variation on chimpanzee cultural behaviour, particularly tool use in the context of foraging (Laland and Janik, 2006; Krutzen et al., 2007). The current consensus is that diversity, distribution, and varying abundance of chimpanzee food sources as well as variation in available tool materials are likely to impact the occurrence, innovation, and maintenance of tool use (Möbius et al. 2008; Schöning et al., 2008; Humle and Matsuzawa, 2002; Gruber et al. 2012; Sanz and Morgan 2013; Gruber et al., 2016; Grund et al. 2019). Two hypotheses: the “necessity”, and “opportunity” hypotheses, relate ecological factors to the emergence of tool use (Fox, Sitompul and Van Schaik, 1999). The necessity hypothesis predicts the emergence of tool use in response to food scarcity (Fox, Sitompul and Van Schaik, 1999), and the opportunity hypothesis proposes that the likelihood for tool use increases when both tool materials and resources requiring tools for exploitation are frequently encountered (Koops, McGrew and Matsuzawa, 2013). Both hypotheses have received empirical support (Koops, McGrew and Matsuzawa, 2013; Yamakoshi, 1998; Sanz and Morgan, 2013; Spagnoletti et al., 2012), and the two hypotheses are not mutually exclusive; thus, both ecological and cultural factors may influence food-related tool use behaviour (Gruber 2013; Grund et al. 2019; see also Rutz and St Clair, 2012).

Ugandan chimpanzees (*Pan troglodytes schweinfurthii*) exhibit a comparatively restricted range of tool use behaviour as compared to other chimpanzee populations and subspecies (McGrew, 2010). For example, the Ngogo and Kanyawara communities in Kibale Forest, and the Sonso community in Budongo Forest (200 km away) respectively show only four, two and one tool use behaviour in relation to food acquisition (Gruber et al., 2012). Nevertheless, the appearance (Hobaiter et al., 2014) and subsequent social spread (Lamon et al., 2017) of the use of a water sponging tool shows that the Sonso chimpanzees are capable of innovation. These observations raise the question of why Ugandan chimpanzees rarely engage in extractive tool use during foraging, despite possessing the cognitive abilities to do so. One possible explanation is that environmental changes have increased food availability (and loss of cyclic food scarcities), causing the Sonso chimpanzees to lose their knowledge of stick-tool manufacture and use in the recent or more distant past (Gruber et al., 2012; Gruber 2013). Humans have played a large role in the forest dynamics and plant species composition of Uganda forests (Reynolds, 2005; Babweteera et al. 2012). For example, four large sawmills were active in Budongo through the 20^th^ century (Synott, 1985), with subsequent logging and species-specific use of arboricides permanently influencing the composition of the forest (Plumptre, 1996), and leading to an increase in the abundance of fruiting trees, such as figs (Tweheyo and Lye, 2003).

While communities living in small riverine fragment forests in close proximity to Budongo (e.g. Bulindi, 60 km away) have been documented to use stick tools during foraging (McLennan, 2011), it is unknown whether the chimpanzees that inhabit the closest large forest between Budongo and Kibale, the Bugoma forest, use tools (Figure 1). In this study we compared the ecological conditions four chimpanzee communities are exposed in two forest areas in order to test hypotheses about the impact of small scale variation in forest ecology on chimpanzee tool using. We compared the potential food availability and tool use opportunities of several communities in the long-term Budongo and newly-established Bugoma forest field sites. We divided our research aim into three parts: firstly, we compared potential chimpanzee food availability in the home range of the respective communities. We expected the diversity and abundance of trees that chimpanzees are known to feed on (Known Feeding Trees) to be highest in the Sonso home range, due to the increased presence of fruit-bearing trees as a by-product of the historical timber production that was centred around this community’s territory (Reynolds, 2005; Gruber, 2013). Furthermore, we expected that the communities’ home ranges differed in Known Feeding Trees species composition.

**Figure 1.**
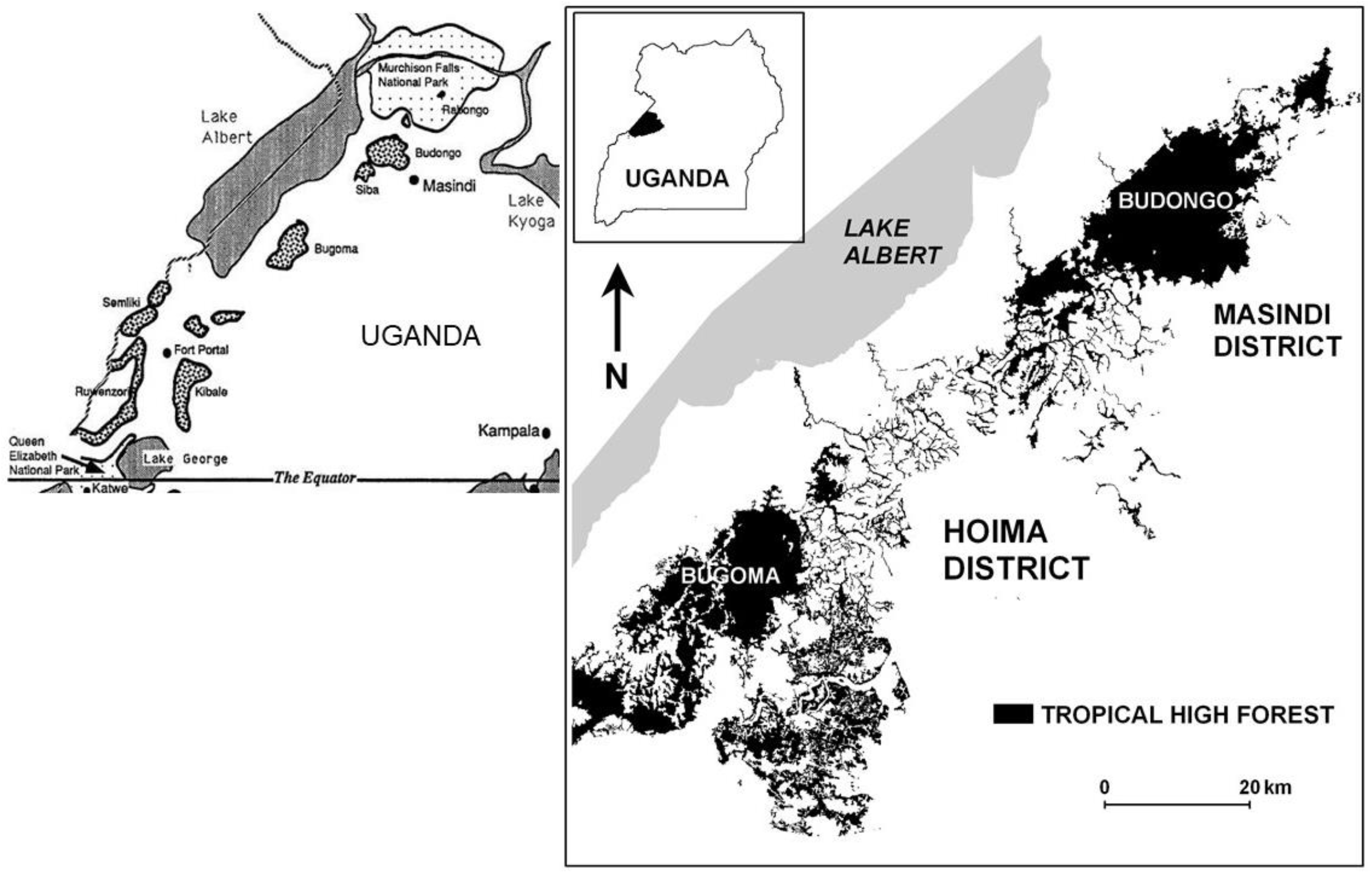
Map of the two study sites: Budongo and Bugoma forest. (Inset map copyright Matthew McLennan; reproduced here with permission)

Secondly, we compared the tool use opportunities in the home ranges of the four communities. We focused on extractive tool use in a foraging context (here the use of sticks to extract insects, as employed by other communities within Uganda, e.g. Watts, 2008). When describing tool use opportunities, we recorded the abundance of termite, *Cubitermes ugandensis*, and ant nests, *Dorylus* spp, and the presence of potential extractive tool materials, such as sticks. The insect species compared were chosen based on previous records of abundance and feeding observations in the Budongo Forest (Hedges and McGrew, 2012). We expected to find fewer tool use opportunities in Sonso as compared to other areas because of the use of poison to control termite populations in recent decades (Reynolds 2005).

Finally, we compared the diet of one chimpanzee community in each forest, Sonso in Budongo and Mwera in Bugoma, using faecal samples from both groups and direct observation of feeding behaviour in Sonso. We expected to find a greater abundance and a more diverse array of seeds in the samples of the Sonso community, due to the predicted difference in Known Feeding Tree abundance described above. To assess the efficacy of faecal analyses, we compared dietary species composition obtained from feeding observations of the Sonso community, with the species composition of their faecal samples. Based on Phillips and McGrew (2013) we expected to identify around 80% of the species from which fruit had been eaten, but only around 20% from which leaves had been eaten, and 60% of species overall in the faecal samples, due to the difficulty of identifying non-frugivory dietary parts at species level (Phillips and McGrew, 2013). We expected the proportional abundance of seeds of a species in the faecal samples to increase the longer we observed the chimpanzees to feed on that species. If faecal analysis in Sonso revealed itself to be a good estimator of diet, we could then use the samples collected in Mwera, to describe the diet of the yet unhabituated chimpanzees.

## METHODS

### a) Study sites

#### i) Budongo Forest

The Budongo Forest is 435km^2^ of continuous semideciduous tropical rain forest, located at the top of the Albertine Rift in western Uganda between 1°37’N-2°03’N and 31°22’-31°46’E with a mean altitude of 1,100m (Plumptre, 1996; Figure 1). Chimpanzee density is estimated to be 1.32 chimpanzees/km^2^ (Plumptre and Cox, 2006). Plumptre (1996) showed that Budongo exhibits a gradual change of tree species composition from the southwest to the northeast, with the southwest having more species associated with Colonizing and Mixed Forest, and the northeast being predominantly *Cynometra* Forest.

#### ii) Bugoma Forest

Bugoma Forest (01°15′N 30°58′E) covers 400km^2^ and is situated between 990 and 1,300 m of elevation (Plumptre, 2010). It is separated from Budongo Forest by around 80km and is the closest major forest to Budongo (Figure 1, Reynolds, 2005), located at an intermediate point between the Budongo and Kibale forests. Much less is known about the history and forest composition of Bugoma than Budongo. In 2006, the density of chimpanzees was estimated to be 1.99 chimpanzees/km^2^ (Plumptre and Cox, 2006).

### b) Study communities

#### Sonso community (Budongo Forest)

The Sonso community (around 70 individuals) has become well-known for its comparatively restricted tool repertoire, particularly in a foraging context (Reynolds, 2005; Gruber et al., 2009). In contrast to all other chimpanzee communities that have been studied long-term, Sonso chimpanzees have never been observed to use sticks to extract food (Whiten et al., 1999). Field experiments involving a hole filled with honey drilled in a log showed that Sonso chimpanzees do not make use of sticks even when put directly into the hole (Gruber et al., 2011). However, Sonso chimpanzees can use objects in a goal-directed manner for water absorption (leaf and moss sponges; Hobaiter et al. 2014; Lamon et al, 2017, 2018), nest building, body care (e.g. leaf-napkin), and social signals (buttress-beat) (Whiten et al., 1999; Gruber et al., 2009; Reynolds, 2005).

#### Waibira community (Budongo Forest)

The Waibira community ranges North-East of the Sonso core area, with overlapping resource use at times. The Waibira community is estimated to total at least 120 individuals; however, as the habituation process started in 2011 some individuals remain only partially habituated. The Waibira chimpanzees have never been observed to use sticks as tools, thus, suggesting the absence of a stick-tool use culture, similar to Sonso. However, since the Waibira chimpanzees have only recently been habituated it is likely that substantial elements of their behaviour and diet remain unknown. They have been observed on two occasions to use leaf-tools to feed on *Dorylus* ‘army-ants’ (Mugisha et al. 2016; Hobaiter, 2019 pers. observation), a task for which other chimpanzee communities employ stick-tools. Insectivory has been previously reported in Budongo chimpanzees but is rare (Newton-Fisher, 1999; Reynolds, 2005) and the Sonso community have not been observed to feed on this species, or to use leaf-tools during insectivory.

#### Kamira community (Budongo Forest)

This community is located North-West of the Sonso chimpanzee range. There is no direct observation of the resident community. As for all other chimpanzee communities in Budongo, there is no direct evidence that the Kamira chimpanzees engage in stick-tool use.

#### Mwera community (Bugoma Forest)

Habituation of the Mwera chimpanzee community in Bugoma began in January 2016. Direct observation of the chimpanzees remains challenging, and their tool repertoire is unknown. A biodiversity survey suggested that Bugoma Forest is ecologically more similar to Kibale than Budongo Forest (Plumptre et al., 2010). In Kibale Forest, the diversity of tree species that produce chimpanzee foods is lower than in Budongo, which has been suggested to impact the increased levels of insectivory and presence of stick-tool use for extractive foraging (Gruber et al., 2012).

### c) Data collection

#### i) Transects

To compare chimpanzee food availability, we conducted 12 500m-long line-transects in both the Sonso and Mwera home range (May-August 2017), and 11 500m-long line-transects in both the Waibira and Kamira home range (July – December 2015; see Supplementary Material for a detailed transect protocol and locations, Table S1-3, Figure S1-2). We identified trees (wherever possible to species level) and ascertained whether it was a Known Feeding Tree based on previous Budongo feeding records from the Sonso and Waibira communities. As fruit represents the major component of chimpanzee diets (Newton-Fisher, 1999), and high fruit abundance has been used to explain the lack of extractive tool use behaviour of the Sonso community (Gruber et al., 2012), we separately compared the abundance and diversity of the subset of Known Feeding Trees from which the fruit is eaten (Known Fruit Trees; Supplementary Material). Occasionally it was not possible to identify the tree; however, this was only ever the case for tree species that chimpanzees were not known to feed on (Non-Feeding Trees). When this occurred, we noted all characteristics as below, but classified the tree as “unidentified”.

The diameter at breast height (DBH) was recorded for all trees that had a DBH of ≥10cm and where at least 50% of the DBH fell within the transect zone. Within Sonso and Mwera community ranges, DBH measurements were recorded for all trees (Known Feeding Trees and Non-Feeding Trees), whereas for Waibira and Kamira, DBH measurements were only recorded for Known Feeding Trees.

When we encountered an insect nest (termite mounds, *Cubitermes ugandensis*, and ant nests, *Dorylus* spp), we took its GPS location, measured its height, determined whether it was active, identified the species (with the help of an experienced field assistant), and assessed surrounding tool availability. Tool availability was assessed by measuring a 5m radius around the mound, scrutinizing a NW (270-360) 90-degree quadrant (or SW (180-270) if NW was not available), and counting all plants capable of producing termite-fishing probes or dipping sticks (classified as twigs, vines, or terrestrial herbaceous vegetation (THV) (Hedges and McGrew, 2012).

#### ii) Feeding observations

The Sonso community numbered around 70 individuals during this study (May through August 2017); however, we only encountered a subset of individuals. We exclusively followed adult chimpanzees, and we balanced as much as possible the choice of focal individuals according to sex and social rank. We followed 8 male and 5 female chimpanzees between 7:30am and 16:30pm. One to six days of feeding observations, with up to a maximum three consecutive days, were made for each focal individual. The total focal sampling time was 246.57 hours (females: mean = 22.29 ± 11.01 hours; males: mean = 15.23 ± 8.93 hours), of which 88.13 hours (females: mean 8.67 ± 5.58 hours; males: mean = 6.92 ± 4.44 hours) were spent feeding by a focal individual. During each follow we used a stop-watch to record the date and time from the beginning of the feeding event until the end. We defined a feeding event as “item placed into mouth, remaining there (or parts thereof) and seen to be either chewed or swallowed” (Phillips and McGrew, 2013). We identified the plant species being eaten (Observed Feeding Plant) with the help of an experienced Budongo Conservation Field Station field assistant. We also noted the food item that was eaten, which we categorized for plant material as: ripe fruit, unripe fruit, mature leaves, young leaves, bark, root, flowers, pith, resin, and rotten wood. We made a subcategory of Observed Feeding Plants from which chimpanzees ate fruit (Observed Fruit Plants, Supplementary Material). Non-plant food items included invertebrates, vertebrates, soil, and honeycomb. We noted the time the focal was lost or out of sight to obtain the total number of minutes a focal was observed for.

#### iii) Faecal samples

In Sonso, we collected samples as soon as possible after a defecation event (never after more than 15min) in a ziplock bag, and noted the producer of the sample, the time, and an estimated percentage of deposited faeces obtained. A sample was considered as complete if ≥95% of the content was collected. Incompleteness was due to faeces consistency, rushed collection due to rapid movement of the focal chimpanzee, or the dispersion of the faeces over a large area (particularly from arboreal defecations).

For the Mwera community, samples were collected opportunistically when found on the ground (only samples estimated to be ≤ 3 days old were collected). We did not record sample completeness for Mwera, as this was not possible to determine without having observed the defecation event. We obtained 105 samples from the Sonso chimpanzees, and 45 from the Mwera chimpanzees. Upon return to the research station, we weighed all samples using a digital scale (Kenex KX digital scale, 400 x 0.1 g), and we added cotton soaked in ethanol into the ziplock bags so that the samples could be stored before processing (up to a maximum of 3-days after collection). All samples were collected between May and August 2017. We processed the faecal samples following McGrew, Marchant and Phillips (2009) in the veterinary laboratory of the Budongo Conservation Field Station (see Supplementary Material for the protocol followed).

### d) Data analysis

We conducted all analyses in R Studio version 1.0.153 (RStudio, 2016).

#### i) Known Feeding Tree abundance

As a considerable proportion of Non-Feeding Trees could not be identified, it was only possible to compare the frequency and diversity of Known Feeding Trees between the ranges of the communities. We compared the abundance of Known Feeding Trees between the communities using Generalized Linear Mixed Models (GLMMs) with forest identity as a random term (random intercept with fixed mean), and chimpanzee community identity and researcher identity as predictor variables.

#### ii) Known Feeding Tree sizes

We log-transformed the DBH data so that they were normally distributed (Figure S3). We fitted GLMMs with forest identity as the random term (random intercept with fixed mean), chimpanzee community identity and researcher identity as predictor variables, and the log-transformed DBH of Known Feeding Trees as the response variable.

#### iii) Overall potential food availability

We used the summed DBH of all Known Feeding Trees of each forest sample as a surrogate measure for potential maximum chimpanzee food abundance. The DBH of a tree is considered a reliable measure of the quantity of fruit it may bear (Chapman et al., 1992). Thus, summing this over all the Known Feeding Trees in a sample yields information on the maximum fruit (and foliage) abundance available to chimpanzees.

We also summed the DBH of all Non-Feeding Trees for Sonso and Mwera, which allowed us to calculate the ratio of the total DBH of Known Feeding Trees to the total DBH of Non-Feeding Trees.

#### iv Diversity of Known Feeding Trees

We chose the Shannon-Wiener index to calculate *α* diversity, as it is the only measure that can be separated into meaningful independent *α* and *β* components when community weights are unequal, and because species are weighted by their relative abundance (Jost, 2007; see Supplementary Material for more information). This weighting means that neither very rare nor very abundant species have a disproportionate impact, and that species richness and species evenness are given equal importance (Jost, 2007; Supplementary Material). We chose the Horn index to calculate *β* diversity, since it uses abundance data (rather than presence-absence data), it relates to Shannon entropy, and when the properties of *β*-metrices were compared, *β*_horn_ scored highly (Barwell, Isaac and Kunin, 2015; Supplementary Material). All diversity indices are entropies (which gives the uncertainty in the species identity of a sample), not diversities, and to be able to interpret them properly we converted them to true *α* and *β* diversities respectively through calculating the exponential of the indices (Jost, 2006; Table S4). We then compared the *α* diversity and the *β* diversity of Known Feeding Trees using GLMMs with forest identity as a random term (random intercept with fixed mean), and chimpanzee community identity and researcher identity as predictor variables. We then calculated the *γ* diversity of each community’s home range (Table S4).

To illustrate our results, we produced a species rank abundance curve and a species accumulation curve using the package “vegan”.

#### v) Known Feeding Tree species composition

We used PRIMER 6 (Clarke and Gorley, 2005) for an Analysis of Similarity (ANOSIM) to determine whether the chimpanzee communities’ home ranges differ in Known Feeding Tree species composition. To visualize dissimilarities between communities we constructed Non-Metric Multidimensional Scaling (NMDS) ordination plots using the package “vegan” in R. Using the package “indicspecies” we looked for the indicator species of each community’s home range and of combinations of two communities (De Caceres, Legendre and Moretti, 2010).

#### vi) Insect nest abundance

We compared the abundance of two species of termite mounds, *Cubitermes ugandensis*, and ant nests, *Dorylus* spp, per transect between the communities’ home ranges using a GLMM with forest identity as the random term (random intercept with fixed mean), and chimpanzee community identity and researcher identity as predictor variables.

#### vii) Abundance of tool material (THV, twigs, and vines)

We contrasted tool availability (i.e. the abundance of THV, vines, and twigs around an insect nest) using a GLMM with chimpanzee community identity and researcher identity as predictor variables, and forest identity as a random term (random intercept with fixed mean).

#### viii) Faecal samples as predictors of chimpanzee diet

As we were not able to distinguish different *Ficus* species in the faecal samples, we combined the *Ficus* species we recorded during feeding observations. Firstly, we fitted a linear model to assess whether the number of plant species identified in the faecal samples could be predicted by the number of Observed Feeding Plants, accounting for the number of hours the respective chimpanzee was observed, and the weight of the faecal sample. If the weight and the hours of observation did not have a significant effect, then we excluded them from the full model.

Secondly, we arcsine square root transformed the proportion of seeds of a given plant species in the faecal samples to meet the assumption of homogeneity of variance (Figure S4). With this as the response variable, we fitted a GLMM with the proportion of time (out of all the recorded feeding time) a chimpanzee was observed to feed on the respective species as the predictor variable, and individual chimpanzee identity as the random term (random intercept with fixed mean). In the full model, we included the total number of hours a chimpanzee was observed, and the total weight of all faecal samples of the respective chimpanzee as covariates. If the reduced models did not differ significantly from the full model, then we excluded the respective covariate from the final model.

#### ix) Abundance and α, β and γ diversity of seeds in faecal samples

We calculated the *α, β* and *γ* diversity of each community from the seeds identified in the faecal samples with the same methods we used when we determined Known Feeding Tree diversity from the transects. However, there were nine samples from Mwera we had to exclude to calculate *β* diversity, because they did not contain any seeds.

After assessing whether the weight of a faecal sample was correlated with the respective predictor variable (using Spearman’s rank correlation), we compared seed abundance and *α* diversity using Wilcoxon tests, and *β* diversity using a two-sample t-test.

## RESULTS

### i) Known Feeding Tree abundance

The communities did not differ in abundance of Known Feeding Trees (GLMM, χ^2^_2,6_ = 2.30, p = 0.32).

### ii) Known Feeding Tree sizes

The mean DBH of Known Feeding Trees differed between the communities’ home ranges (GLMM, χ^2^_2,6_ = 7.86, p < 0.020). Only Sonso differed from all other communities, with a greater mean DBH (Table 1, Table S6).

**Table 1.**
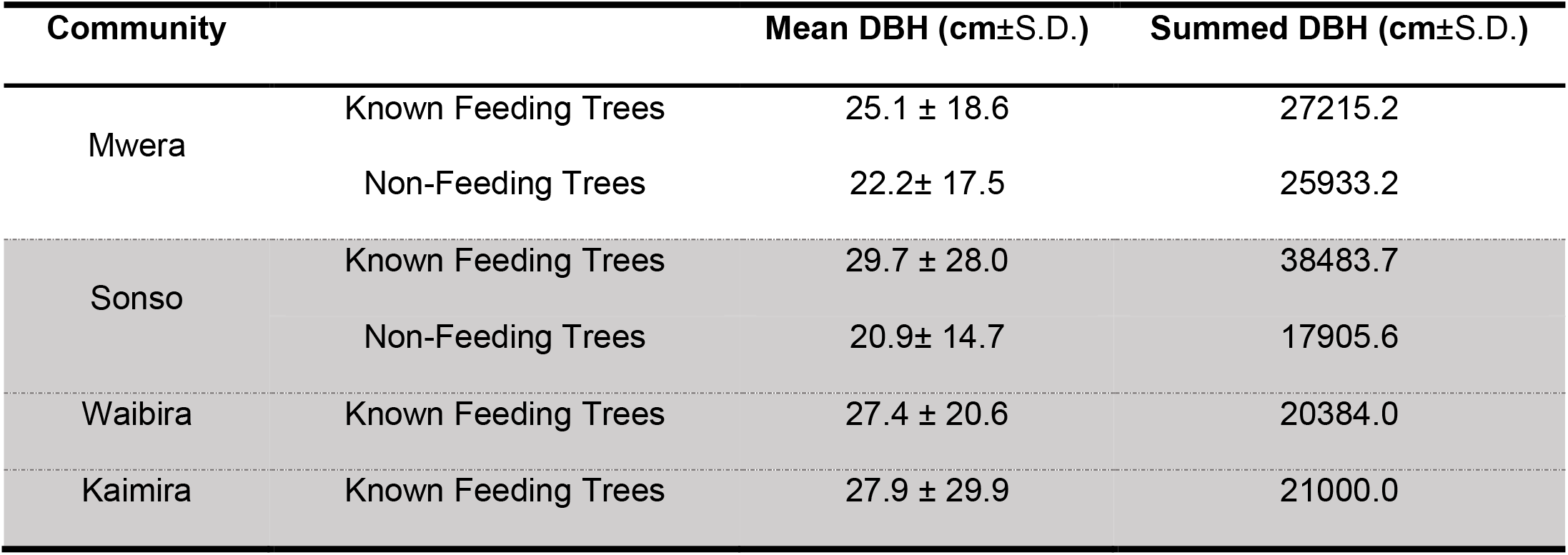
The mean DBH and total DBH of all trees encountered during the transects (for Known Feeding Trees and Non-Feeding Trees) for each community. The communities from the Budongo forest are shaded grey.

### iii) Potential food availability in each community’s home range

The summed DBH of Known Feeding Trees was highest for Sonso, which was almost twice as high as the summed DBH of the other two Budongo Forest communities: Waibira and Kamira. Mwera in Bugoma Forest had the second highest summed-DBH value. For Sonso, the summed DBH of Known Feeding Trees was over twice as high as the summed DBH of Non-Feeding Trees; the ratio was roughly 1:1 for Mwera (Table 1).

### iv) Diversity of Known Feeding Trees

The *α* diversity of Known Feeding Trees differed between the communities’ home ranges (GLMM, χ^2^_2,6_ = 11.97, p < 0.001), where the *α* diversity of Sonso was roughly 2.5 times greater than the *α* diversity of any other community (Table 2, Figure 2). Sonso was the only community that had a greater *α* diversity of Known Feeding Trees than any other community (Table S7). The *β* diversity of Known Feeding Trees differed between the communities’ home ranges (GLMM, χ^2^_2,6_ = 8.55, p = 0.014), where Kamira had a lower diversity than any other community, albeit not strongly (Table S7). The *γ* diversity of Known Feeding Trees in Sonso was roughly 2.60 times greater than in any other community (Table 2, Figure 2).

**Figure 2.**
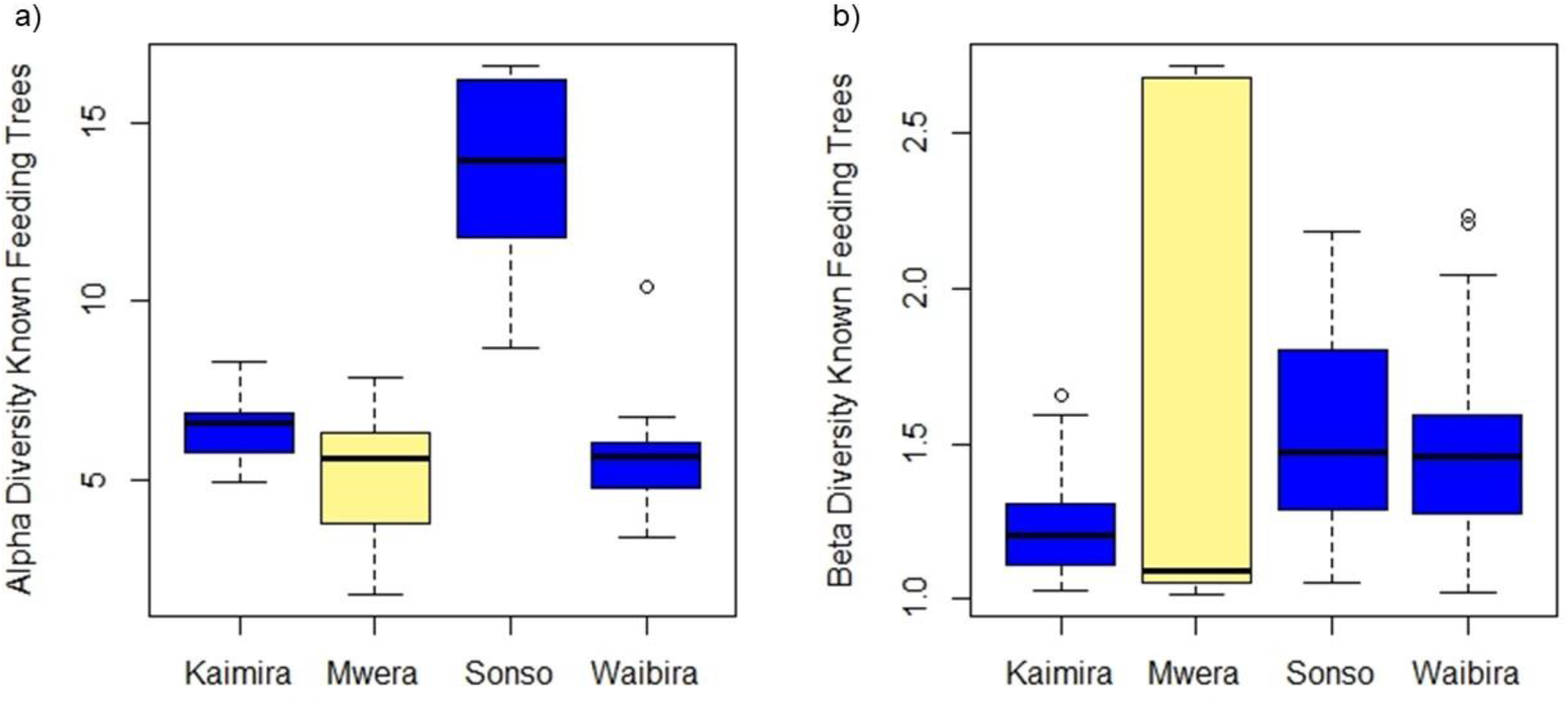
Boxplot illustrating the median and the upper and lower quartiles of the (a) alpha and (b) beta diversity of Known Feeding Trees of the home ranges of the four communities. The communities from Budongo forest are shaded in yellow, and the Mwera community (Bugoma forest) is shaded in blue.

**Table 2.**
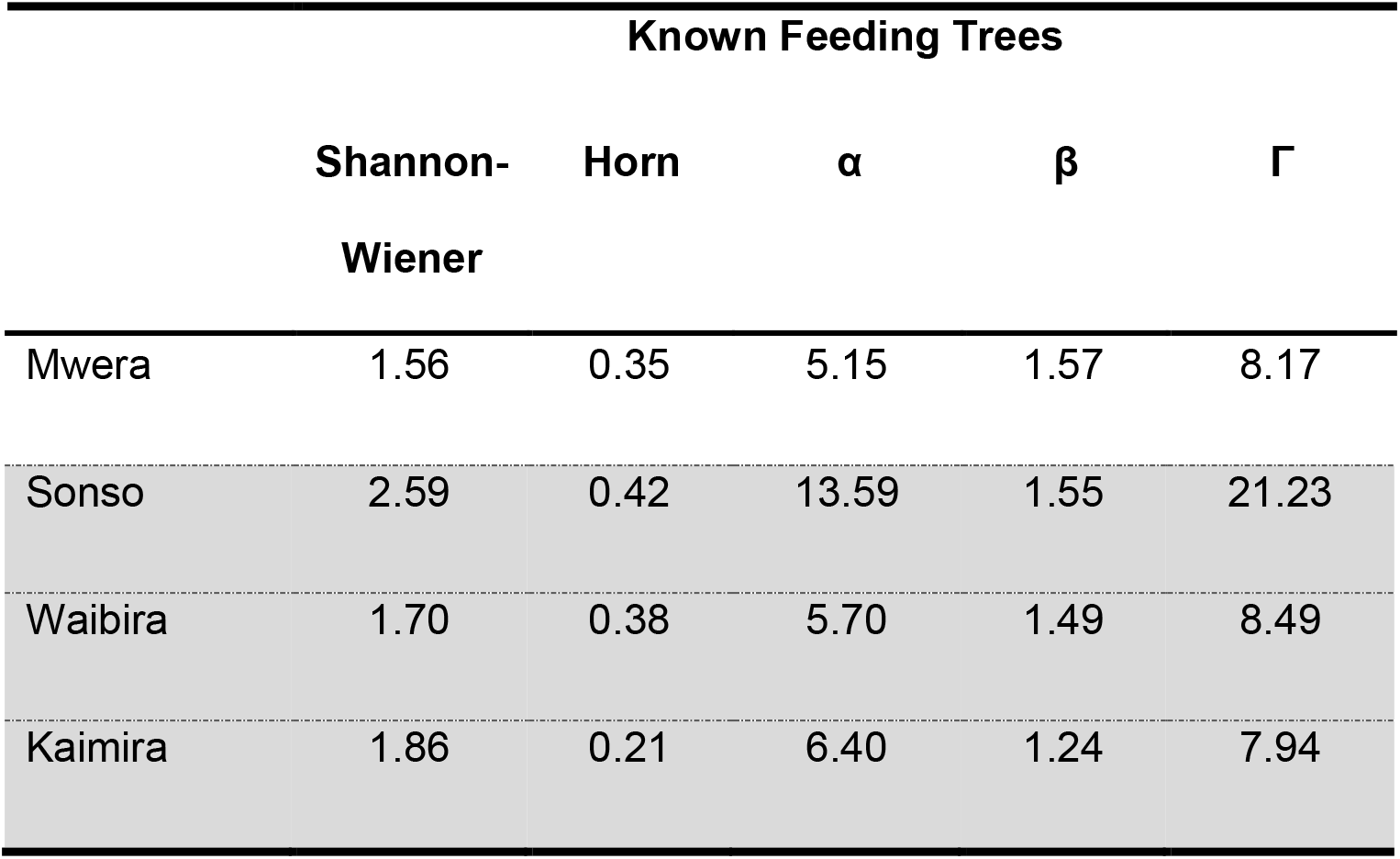
Diversity indices and Alpha, Beta and Gamma diversity of Known Feeding Trees for each community’s range. The communities from the Budongo forest are shaded grey.

The abundance of species, represented by Known Feeding Tree species richness and species evenness, were greatest for the Sonso community’s range (Figure 3a). The species accumulation curves are in line with the results from the diversity analyses since the number of Known Feeding Tree species recorded was highest for Sonso for a given number of transects (Figure 3b).

**Figure 3.**
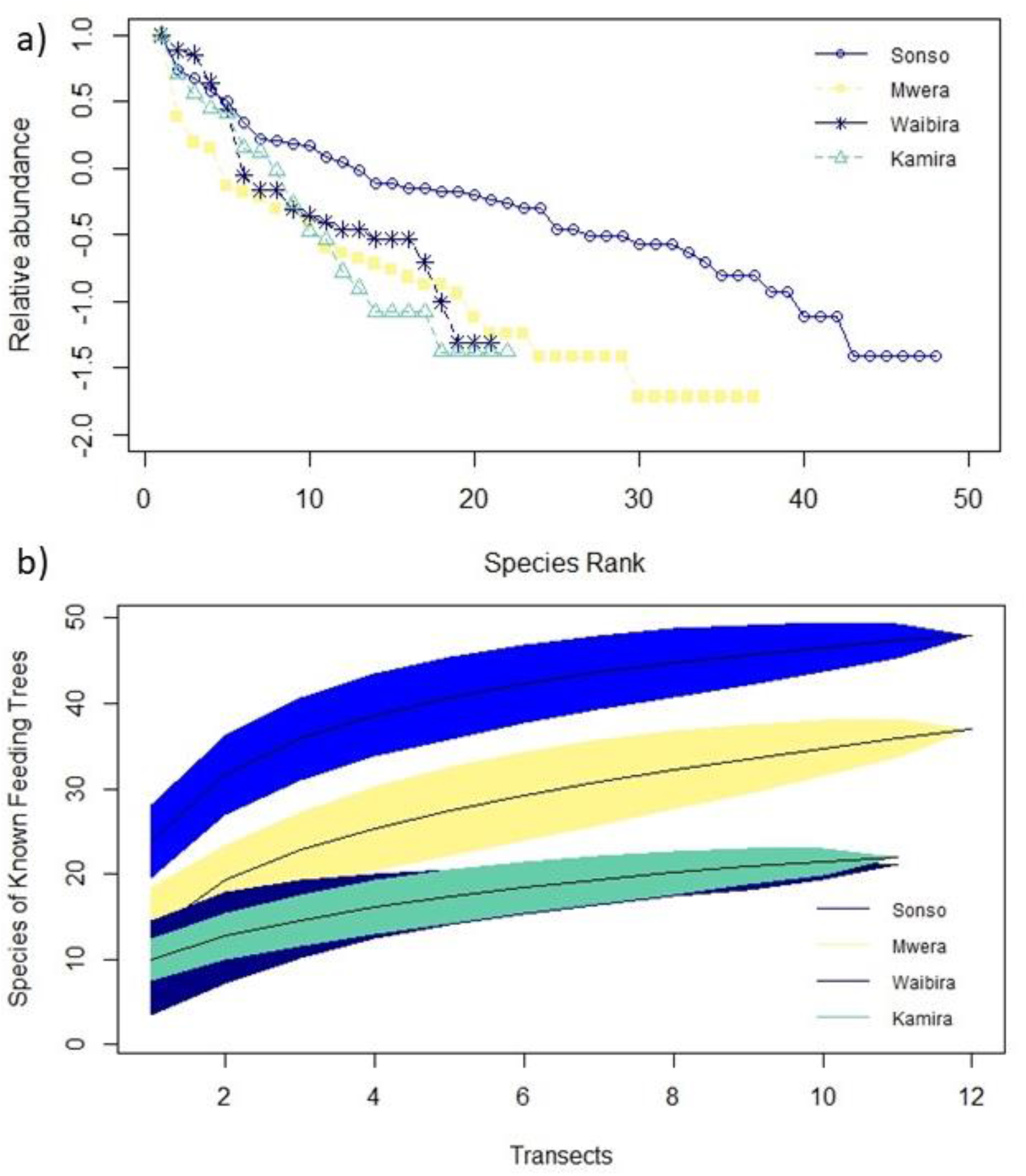
(a) Species rank abundance curve of Known Feeding Tree species recorded during the transects for each community’s home range. Species richness is illustrated by the length of the line (i.e. the number of species) and species evenness is indicated by the steepness of the line (i.e. the abundance of a species in relation to the other species in the forest), which are both highest for Sonso. (b) Species accumulation curve for Known Feeding Trees depicting how the number of recorded Known Feeding Tree species increases the more transects we conducted. For any given number of transects, the number of recorded species was highest for Sonso, followed by Mwera. The areas shaded in show the 95% confidence intervals.

### v) Known Feeding Tree species composition

The Known Feeding Tree species composition differed between the community home ranges (ANOSIM, R= 0.41, p= 0.001). The home ranges of Waibira and Kamira had a similar species composition and relative abundance of Known Feeding Trees, whereas Sonso and Mwera differed in this respect (Figure 4).

**Figure 4.**
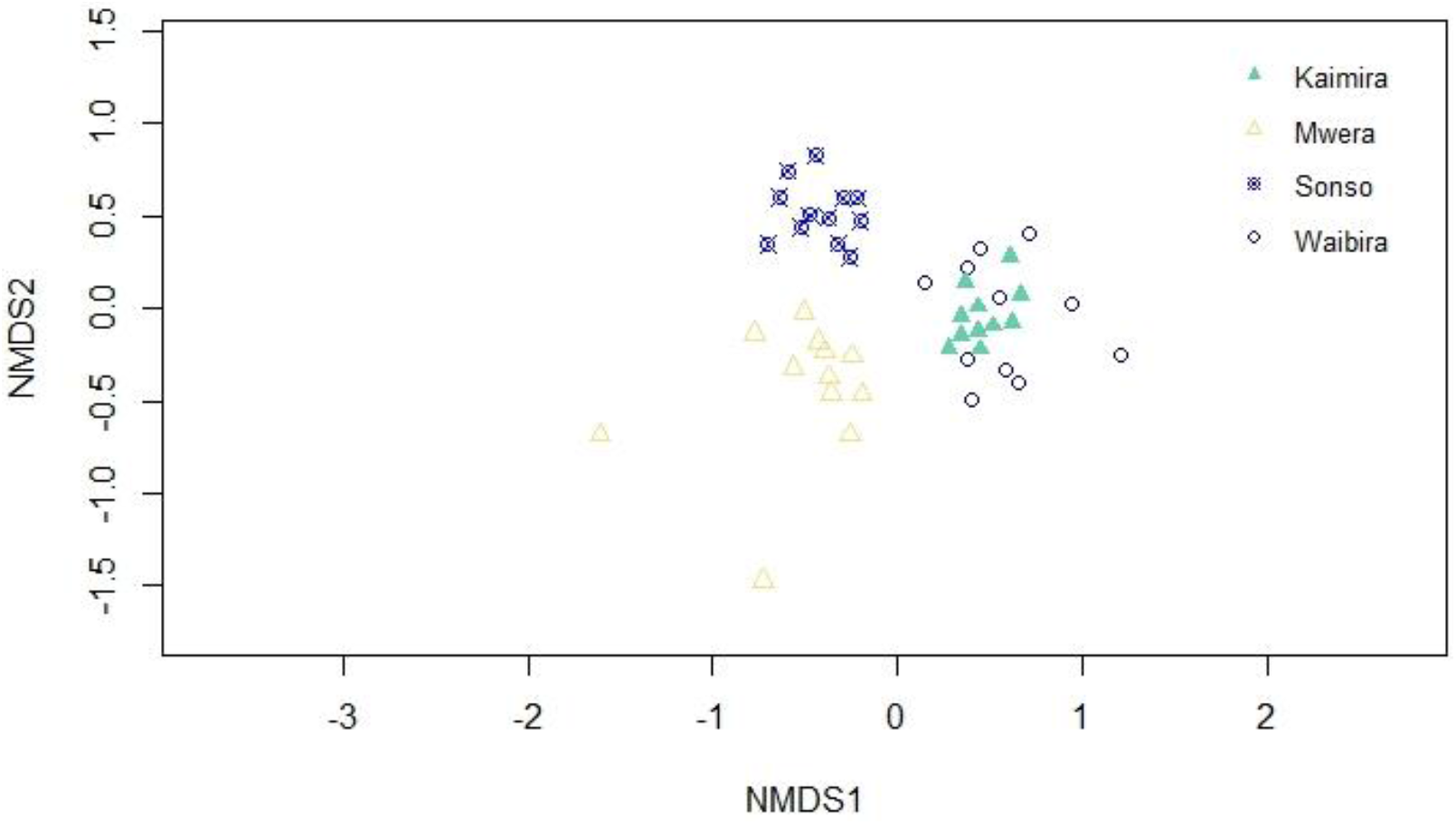
Non-Metric Multidimensional Scaling ordination plot of dissimilarities between the Known Feeding Tree community of the home ranges of the different chimpanzee communities. Each point represents a different transect and the closer the points are the more similar the transects are to each other. Thus, points of a given community are clustered together. Kaimira and Waibira are very similar to each other, whereas Sonso and Mwera differ, with more pronounced variation between the transects within Mwera.

We only found indicator Known Feeding Tree species for the home ranges of Sonso and Mwera, implying that Kamira and Waibira home ranges are populated by species that are commonly found at the other communities’ home ranges (Table S8). We found ten indicator species for Sonso (all indigenous rather than introduced species), where *Trichilia rubescens, Teclea nobilis*, and *Croton sylvaticus* had the highest indicator values, meaning that they are the most characteristic species. Mwera was characterized by three species: *Morus lactea, Chrysophyllum muerense*, and *Sterculia dawei* (also indigenous species). We found three community combinations with indicator species, all of which included Sonso. Thus, the home range of Sonso appears to contain many Known Feeding Tree species that are absent or rare at other home ranges.

### vi) Insect nest abundance

The abundance of termite and ant nests differed between the forests (GLMM, χ^2^_2,6_ = 7.18, p = 0.028), where Kamira had a greater abundance than Mwera and Sonso, and Waibira a greater abundance than Sonso (Figure 5, Table S10).

**Figure 5.**
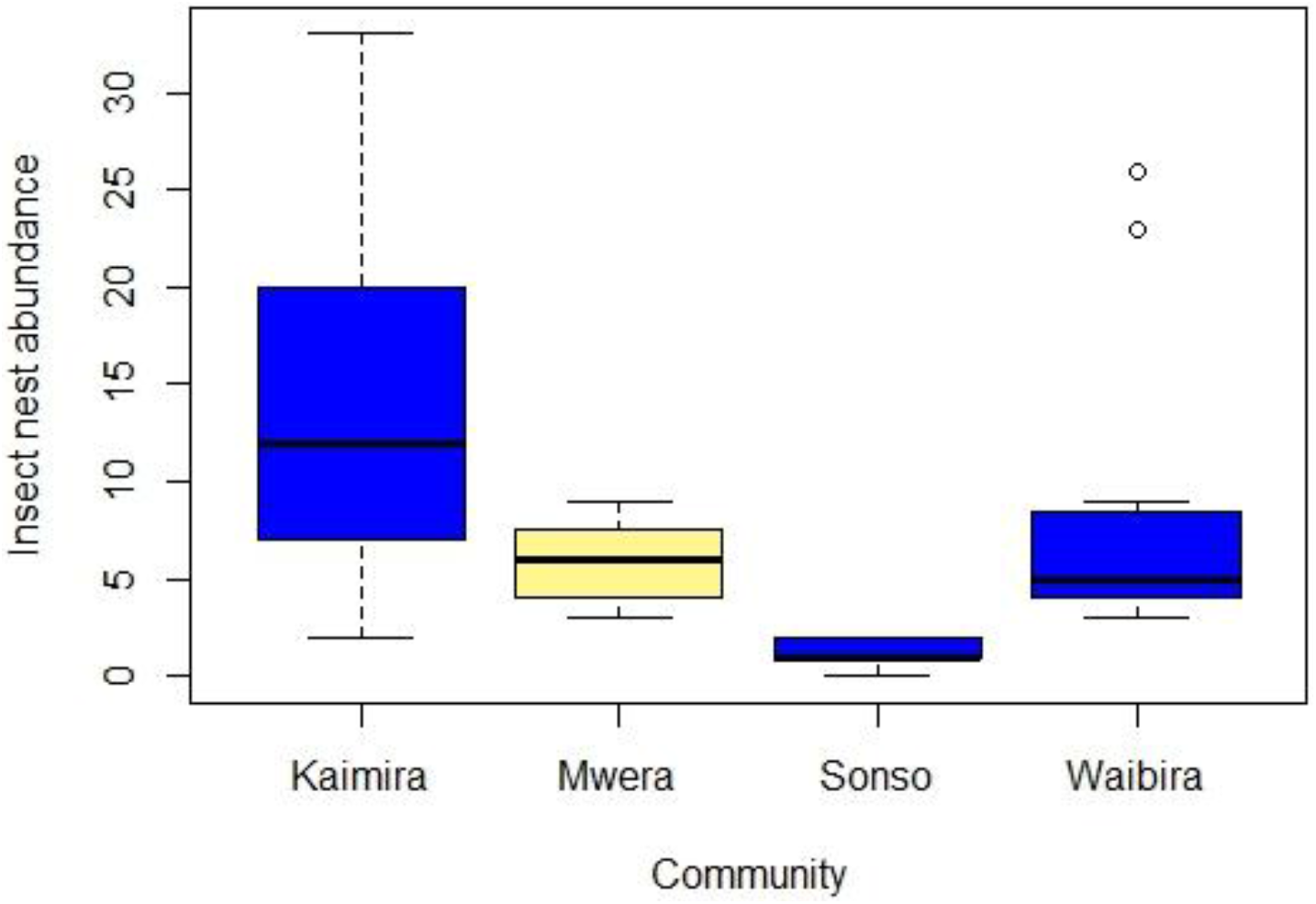
Boxplot illustrating the median and the upper and lower quartiles of the number of termite mounds and ant nests per transect for each community’s home range. The communities that are found in Budongo forest are shaded in yellow.

### vii) Abundance of tool material (THV, twigs, and vines)

The abundance of THV, twigs and vines did not differ between the communities’ home ranges (GLMM, *X*_2,6_^2^ = 4.88, p = 0.087).

### viii) Faecal samples’ predictive power

Around half (48.73%) of the Observed Plant Species in Sonso were found in the faecal samples. Fig species accounted for 79.53% of the seeds, followed by *Broussonetia papyrifera* (11.24%) and *Psidium guajava* (8.76%; Table 3). From the feeding observations, figs were most heavily fed on, accounting for 30.82% of the feeding time. Similar to the faecal samples, *Broussonetia papyrifera* was an important feeding species (17.10%; Table 3), but *Psidium guajava* only accounted for 2.59% of the feeding time. In contrast, we observed the Sonso chimpanzees to feed on *Cordia millenii* 17.92% of the time (Table 3), but just 0.039% of the seeds found in faecal samples were of that species (Table 3). Overall, species richness and evenness appear to be lower for the faecal samples.

**Table 3.**
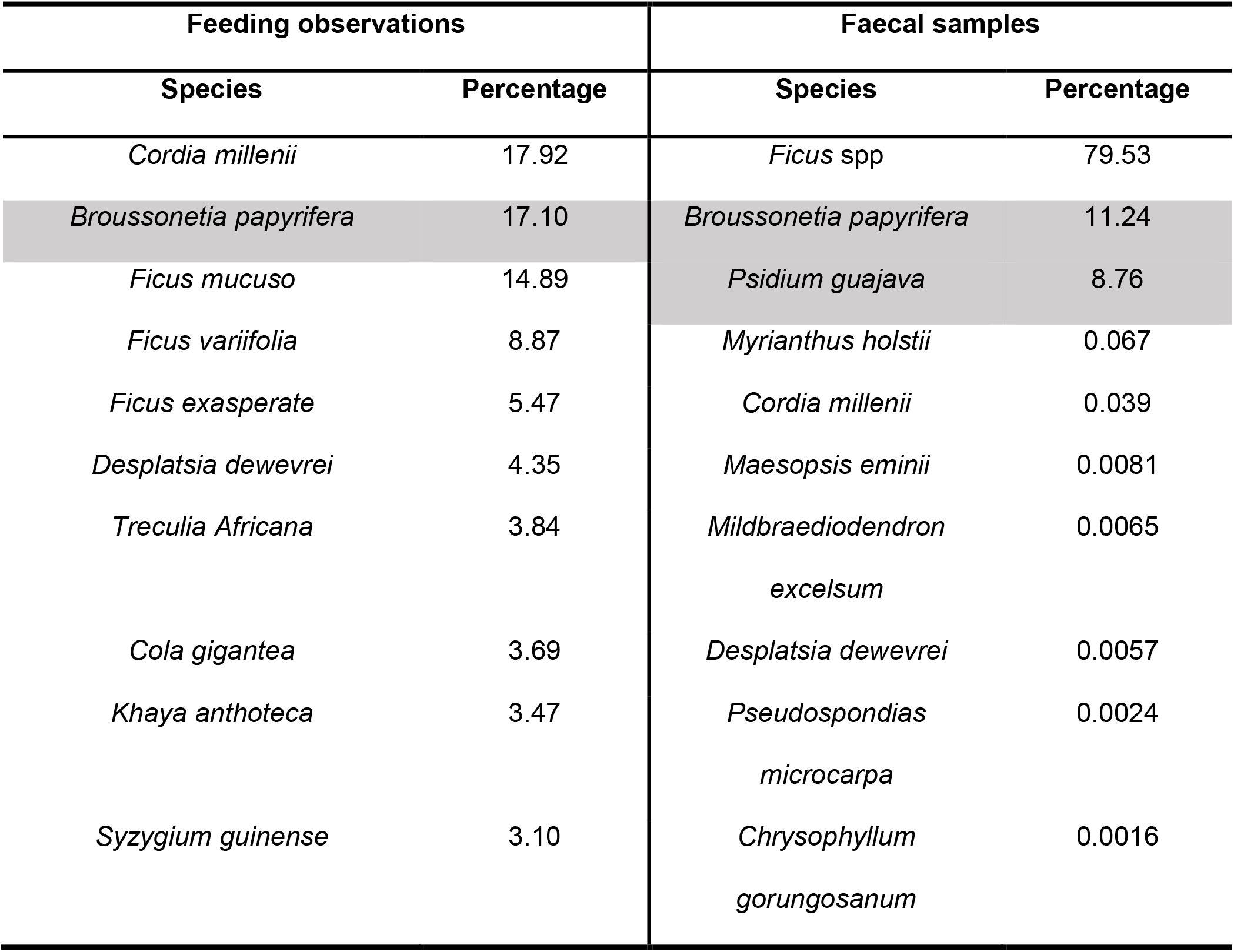
Preferred Sonso chimpanzee feeding trees described by the percentage of time spent feeding. The species shaded in grey are non-native species introduced to the forest.

The number of plant species found in the faecal samples was not affected by the number of Observed Feeding Plants (lm, F_1,11_ = 2.03, p = 0.18), when the weight of the faecal samples, and the duration of observation were excluded due to their non-significance (lm, F_1,10_ = 0.73, p = 0.42; lm, F_1,10_ = 4.85, p = 0.055; Figure S5).

The total weight of the faecal samples for a given chimpanzee did not affect the proportion of a particular species in the faecal samples (GLMM, χ^2^_1,6_ = 0.040, p = 0.84) and was thus excluded from the model, as were the hours a chimpanzee was observed for (GLMM, χ^2^_1,6_ = 0.028, p = 0.87). The greater the proportion of time a chimpanzee fed on a plant species, the higher the proportion of its seeds in the faecal samples (GLMM, χ^2^_1,4_ = 161.9, p < 0.0005, coefficient = 0.83; Figure S6).

### ix) Abundance and α, β and γ diversity of seeds in faecal samples

Seed abundance positively correlated with the weight of the sample (Spearman’s rank correlation, S_151_ = 426640, p = 0.0029). However, the average weight of a sample did not differ between the Sonso and Mwera communities (two sample *t*-test, *t*_149_ = 1.032, p = 0.30), meaning that we were able to exclude weight from subsequent analyses. The mean abundance of seeds per faecal sample was greater for the Sonso community (2060.82 ± 366.99 s.e.) than the Mwera community (839.46 ± 441.28 s.e.; Wilcoxon, W_151_ = 1052, p < 0.001; Figure S7).

Given that the data of the *α* diversity of seeds of faecal samples did not exhibit homogeneity of variance, we were unable to perform an ANCOVA. However, the weight of a faecal sample did not correlate with the *α* diversity of that sample (Spearman’s rank correlation, S_151_ = 571800, p = 0.97), and was thus excluded from the model. The *α* diversity of seeds in faecal samples was 1.5 times greater for the Sonso community (Wilcoxon, W_151_ = 518, p < 0.001; Table 4; Figure S8).

**Table 4.** Diversity indices, and Alpha, Beta and Gamma diversity of the seeds found in the faecal samples of the two communities.

| Sonso | Mwera |
| --- | --- |

| Shannon-Wiener | Horn | $\alpha$ | $\beta$ | $\gamma$ | Shannon-Wiener | Horn | $\alpha$ | $\beta$ | $\gamma$ |
| --- | --- | --- | --- | --- | --- | --- | --- | --- | --- |
| 0.43 | 0.46 | 1.61 | 1.70 | 2.42 | 0.054 | 0.50 | 1.07 | 1.85 | 1.75 |

Due to the nature of how *β*_horn_ is calculated, it was not possible to assess whether the weight of the sample had an effect. However, as shown above, weight did not appear to differ between the two communities. The *β* diversity was 1.09 times greater for the Mwera community (two sample t-test, *t*_5949_= −5.34, p < 0.001; Table 4; Figure S8), but γ diversity was 1.38X greater for the Sonso community (Table 4).

The species rank abundance curve of the species identified from seeds in the faecal samples suggests a lower species evenness and species richness in the diet of the Mwera chimpanzees (Figure 6a). The species accumulation curve suggests a greater species richness in the diet of the Sonso chimpanzees (since for a given number of faecal samples, the number of species recorded is greater for Sonso), although there is considerable overlap of the 95% confidence intervals (Figure 6b).

**Figure 6.**
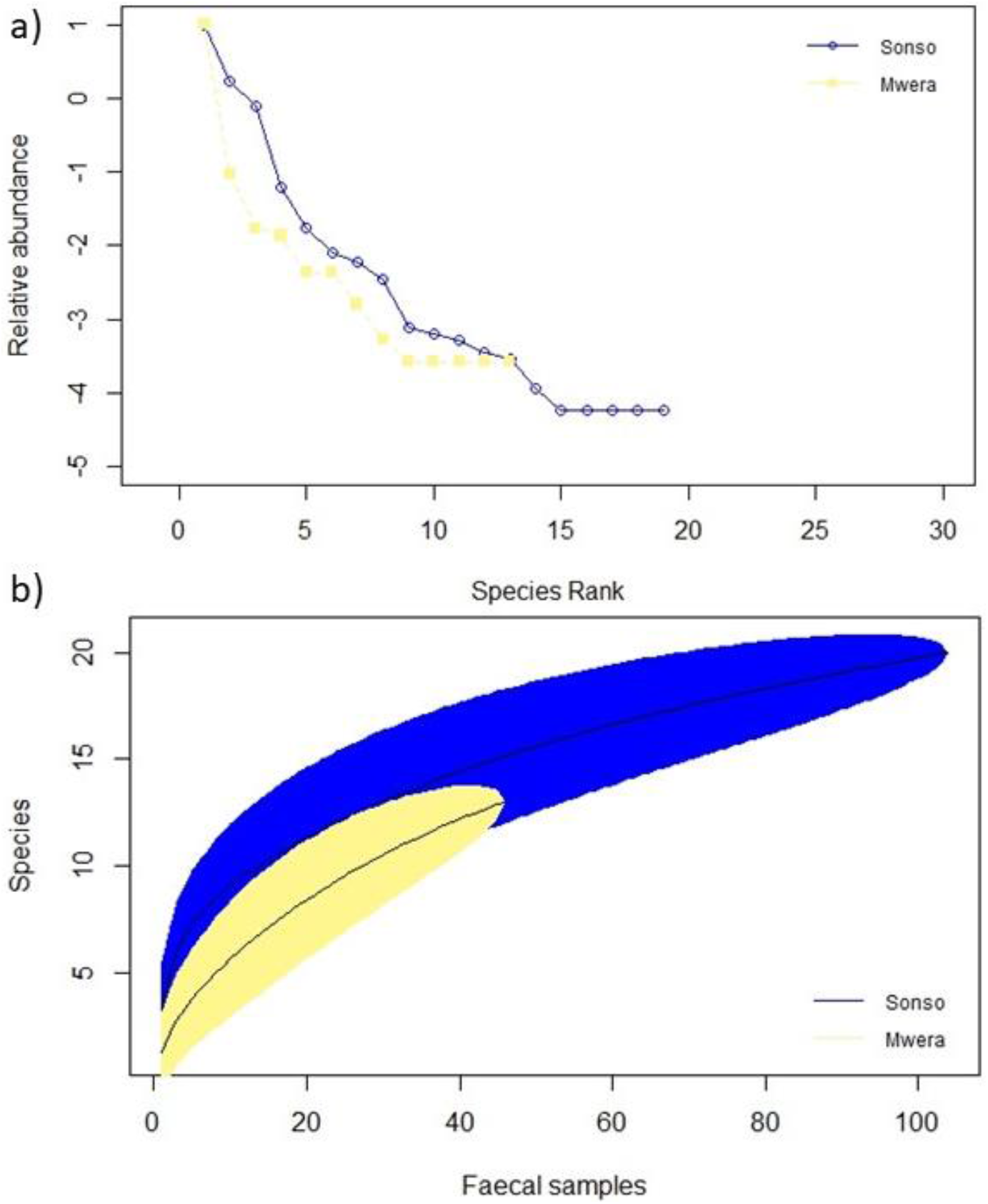
(a) Species rank abundance curves for the seeds identified in the faecal samples for the Sonso and Mwera chimpanzee communities. Species richness is illustrated by the length of the line (i.e. the number of species) and species evenness is indicated by the steepness of the line (i.e. the abundance of a species in relation to the other species in the forest): both are higher for the Sonso chimpanzees. (b) Species accumulation curve, illustrating how the number of species identified from seeds increases with the number of faecal samples analysed for both chimpanzee communities, which is higher for Sonso. The areas shaded in represent the 95% confidence intervals, which overlap.

## DISCUSSION

We analysed a range of ecological indicators that might influence the likelihood of wild chimpanzees engaging in extractive tool use in two Ugandan forests. We found that, while most groups’ territories did not differ substantially in these factors, the Sonso community’s home range in the Budongo forest had both a higher diversity and biomass of chimpanzee feeding species. In addition, Sonso chimpanzees had the greatest number of Known Feeding Tree species unique to their range, and the lowest termite and ant nest abundance. The abundance of potential tool materials did not differ between any of the communities’ home ranges. Faecal samples underestimate food species richness, under-representing some species, and over-representing others. However, they nevertheless provided valuable insight into chimpanzee diets and remain a useful tool for describing unhabituated chimpanzee feeding behaviour. The greater abundance and diversity of seeds we found in the Sonso chimpanzees’ faecal samples is likely to reflect a genuine difference in diet between the Sonso and Mwera chimpanzees, which may have consequences for their tool use behaviour.

### Potential food availability differs between chimpanzee communities’ home ranges

While the abundance of chimpanzee feeding trees did not differ between the communities’ home ranges (either within or between forests); there was a small difference in biomass – a measure of the potential fruit abundance – suggesting, that the Sonso community benefits from particularly high food availability. Systematic managed logging occurred across the Ugandan National Forest Reserves; however, within them some areas were designated ‘Nature Reserves’ with no legal logging taking place. Only the Kamira chimpanzee’s home range overlapped with one of these reserve areas, making it hard to draw meaningful inferences about the impact of managed logging. However, the presence of widespread illegal logging may better explain the observed pattern of variation in biomass. Illegal loggers particularly target large mature trees. While illegal logging occurs across all four communities’ home ranges, the presence of an active research station in the Sonso community’s range for almost 30-years appears to have conferred significant protection (Babweteera et al., 2012), allowing a greater proportion of trees to mature.

The *α* diversity of Known Feeding Trees was greatest for Sonso, roughly 2.5X greater than any other community. This difference means that at the local scale (i.e. per transect) Sonso had the greatest diversity. The *β* diversity of Known Feeding Trees was lowest for Kamira, possibly because its location near a nature reserve may mean that the area is relatively uniform (i.e. a low compositional turnover between transects) in climax species. However, the difference in diversity between areas was quite small. As predicted, the total diversity seemed to be greatest for Sonso (supported by the highest γ diversity). The results are in line with previous studies which found that, when compared to chimpanzees in Kibale, the Sonso chimpanzees diet contains a higher diversity of food items which may limit any negative effect of seasonal food shortages and reduce the necessity to use tools to extract alternative food resources (Reynolds 2005; Gruber et al. 2012). Nevertheless, a cross-seasonal survey of actual chimpanzee food availability is needed to strengthen these arguments.

Many indicator species were found for Sonso (alone or in combination with another community’s home range). Thus, the Sonso chimpanzees’ home range has many species that are absent or rare in other communities’ home ranges. This diversity further highlights the range of feeding options available to Sonso chimpanzees, again potentially reducing the impact of cyclic food scarcities in more widely available feeding species.

Does the higher potential food availability in Sonso sustain greater chimpanzee and other potential competitor species densities (such as other frugivorous primates and birds)? A similar density of small mammals and birds in Budongo and Bugoma suggests that this is not the case (Owiunji, 2000; Plumptre et al., 2010), and the density of chimpanzees is slightly lower for Budongo, suggesting similar levels of intraspecific and interspecific competition across the forest areas (Plumptre and Cox, 2006).

### Dietary differences between Sonso and Mwera

Based on previous studies of chimpanzee faecal samples (Phillips and McGrew, 2013) we expected to identify around 60% of the species that we had observed the Sonso chimpanzees feeding on in their faecal samples, but we were only able to identify around half. Faecal samples appear to represent the frugivorous component of the Sonso chimpanzees’ diet relatively well, but not the folivorous component and we were unable to identify any species from the leaf fragments found in the faecal samples.

A comparison of the most common species in the faecal samples to the most frequently recorded Observed Feeding Plants, indicated that, whilst there is considerable overlap, fruits that produce many small seeds (such as *Broussonetia papyrifera, Ficus* spp., and *Psidium guajava*) are overrepresented in the faecal samples. In contrast, fruit that produce large seeds, from which the chimpanzees scrape off the flesh with their teeth but then rarely swallow the seeds, are underrepresented. This bias likely explains our observations for *Cordia millenii*, the species we recorded as the one chimpanzees spent the most time feeding on, but which was rarely found in their faecal samples. However, as predicted, the greater the proportion of time chimpanzees fed on a species, the higher the proportion of its seeds were found in the faecal samples. Within this model we aimed to go beyond the discriptive results of previous studies (e.g. Phillips and McGrew, 2013), but it proved challenging due to zero-inflation. The zero-inflation was probably due to us being unable to identify plant species in the faecal samples from which only leaves had been eaten, and chimpanzees feeding on fruits whose large seeds were not regularly swallowed. The zero-inflation of the model is problematic, but it will be difficult to avoid in future studies, even with a larger sample size, given the number of feeding species that are not well represented by or easily identified in faecal samples.

As predicted, the abundance and *α* diversity of seeds were greater for Sonso chimpanzees’ faecal samples as compared to the Mwera samples, suggesting that chimpanzees in the Sonso community have a more diverse diet than the latter. The *β* diversity of seeds was slightly greater for the Mwera chimpanzees’ samples, however, we were unable to control for individual identity in sample collection (and may have collected disproportion numbers of samples from some individuals, particularly large mature males who are easier to find and approach in less well habituated groups). Furthermore, the sample size for Mwera was roughly half that of Sonso, and to calculate *β* diversity we had to exclude samples from which we did not record any seeds (which only occurred in Mwera samples). Both factors may have artificially increased the estimate of *β* diversity for Mwera samples and thus the estimates are not sufficiently reliable to warrant interpretation.

Nevertheless, across measures, faecal samples provided a relatively good insight into the identity of the most important feeding species, and remain an important tool in the description of chimpanzee feeding behaviour, particularly for comparison of dietary differences between communities.

### Variation in extractive tool use opportunities between Budongo and Bugoma Forests

Insectivory is rare in Budongo as well as in Kibale forest chimpanzees (Watts et al., 2012; but see Mugisha et al. 2016), but common in other mid-western areas of Uganda (Semliki: Webster et al., 2014; Bulindi: McLennan, 2014; Kalinzu: Koops et al., 2015). Across the four communities and two forests in our sample, Kamira had the greatest abundance of termite mounds, *Cubitermes ugandensis*, and ant nests, *Dorylus* spp, and Sonso the lowest, both in Budongo. The presence of the research station at the centre of the Sonso community territory likely afforded significant protection from illegal logging of mature trees over the past 30-years. However, Sonso also experienced the highest levels of active forest management during the decades of timber production. This management appears to have changed the composition of tree species (as seen in the diversity of species available for chimpanzee feeding) and included the active use of tree-species pesticides and termite mound poisoning (Reynolds 2005). In contrast, the Kamira chimpanzees’ territory overlaps an area designated as a Nature Reserve during timber production, and likely received the least invasive use of management practice. The variation in insect nests may reflect the variation in the pattern of human impact, with active management for timber production a possible factor in explaining the low abundance of insect nests even decades after production stopped.

While this suggests that the Sonso chimpanzees have the lowest opportunity to feed on termites and ants within our sample, opportunity is unlikely to fully explain the absence of extractive tool use during foraging in Sonso chimpanzees (Grund et al. 2019). The abundance of tool materials did not differ between the communities’ home ranges, and, while low, the abundance of termite and ant nests in Sonso was previously found to be within the range of the densities at sites were tools are used to extract insects (Hedges and McGrew, 2012).

### Potential implications of the measured ecological variables on tool use

Our study suggests that a high diversity and biomass of Known Feeding Trees in Sonso underscores the Sonso chimpanzees’ comparatively diverse and fruit-rich diet, and supports previous work suggesting that the Sonso chimpanzees have the most diverse food availability out of five Ugandan forest locations (Gruber et al., 2012). In previous work the Kanyawara chimpanzees in Kibale forest were suggested to live in the least favourable environment in terms of food diversity and quality but did not exhibit more extensive tool use than other Uganda chimpanzee communities. As a result, the authors concluded that ecological conditions could not completely explain observed differences in extractive tool use for foraging (Gruber et al., 2012). In this study, we again find differences in food diversity and quality across communities, even those with adjacent territories within the same forest area. However, it is not known whether Kamira and Mwera chimpanzees engage in extractive tool use behaviour (beyond sponging for water), and it appears that Waibira chimpanzees, like Sonso chimpanzees, do not use stick tools (Mugisha et al., 2016), albeit after only a more limited number of years of observation (currently 8 yrs). Nevertheless, the observations available suggest that chimpanzees in Waibira have developed additional strategies relying on tool use to acquire valuable proteins (using leaf tools to feed on *Dorylus* ants, Mugisha et al., 2016) compared to the Sonso chimpanzees, where this behaviour has never been observed in 25 years of continuous study. Once the four communities’ tool repertoire is known, it can be mapped together with our understanding of current and historic socio-ecological conditions.

The study of chimpanzees in forest areas that lie intermediate between Budongo where stick-tool use appears absent, and forest areas with communities that do employ stick-tool use (for example in Kibale) is of significant interest. The Bulindi chimpanzee community lives in a small riverine fragment less than 60km from Budongo, and has been shown to employ stick-tool use to obtain honey (McLennan, 2014). The Budongo and Bulindi communities show dramatic differences in habitat, particularly in respect to chimpanzee feeding tree species, which may underlie their variation in diet and tool use (McLennan, 2014). But it remains unknown whether or not the Bulindi chimpanzees a) re-innovated tool use in response to the degradation of their fragmenting habitat, b) differed from other Budongo communities before their forest areas diverged, or c) retained extractive stick tool use while other Budongo communities lost it. The difficulty that chimpanzees experience in re-innovating tool use (e.g. Gruber et al., 2011) suggests that option a) is either unlikely, or requires substantial pressure. Once more information on the tool repertoires of the four communities we studied becomes known, the ecological data we collected will help answer these questions.

Nevertheless, based on our ecological findings, we predict that the Mwera community engages in extractive tool use behaviour in a foraging context. Potential food availability and tool use opportunities were comparatively low within the Mwera chimpanzee range, they appear to have a less diverse diet than the Sonso chimpanzees, and are geographically closer to the chimpanzee communities in both Kibale and Bulindi that use stick-tools. As habituation improves, direct observations will allow us to test this prediction.

Future work exploring tool use, and variation in tool-using across chimpanzee communities in the Budongo and Bugoma forests will be of particular interest to explanations for its unusual absence in the Sonso chimpanzees. The information obtained can then be paired with the ecological and dietary insights we gained in this study and contribute to our wider understanding of the interactions between ecological and cognitive aspects of chimpanzee tool use.

## ETHNICAL APPROVAL

All applicable international, national, and/or institutional guidelines for the care and use of animals were followed.

## INFORMED CONSTENT

The research did not involve human participants. Research on wild chimpanzees was in accordance with international guidelines and that of the research station.

## DATA AVAILABILITY

All data was collected by the authors. If any raw data and/or analyses (incl. R-code) are wanted the corresponding author, Iris Berger, can provide them.

## ACKNOWLEDGEMENTS

We thank the National Geographic Society and the Davis Expedition Fund for their generous financial support. We thank Helene Maurer, Gideon Monday, and Adaku Amon for their help with the data collection, and Dr Matthew McLennan for help with seed identification and permission to use his map of Uganda.

