## Supplementary Material for "Ecological and dietary differences between Ugandan chimpanzee communities with possible implications on tool use"

### Transect locations

The location of the transects in the Sonso range was based on a previous study (Hedges and McGrew, 2012), where 12 line-transects of 6m width and 500m length were conducted along the existing grid system of trails. Six transects ran north-south, and six ran east-west, with the sites of the transects balanced for the two main forest types: Mixed and Swamp. Since conducting transects along trails may have confounded our results (as no trails exist in Bugoma), we walked 50m east into the forest from north-south trails, before conducting a transect parallel to the trail. For east-west trails, we walked 50m north before conducting a parallel transect (using Garmin BaseCamp) so that all transects were within forest compartments (Supplementary Table 1, Supplementary Figure 1). The start point for north-south transects was always north, and for east-west transects was always east.

We used the same transect arrangement (i.e. the same relative position of the transects) for the Mwera range, where the exact location was based on chimpanzee range data from January 2016 to April 2017, i.e. we attempted to place the transects in a way that best overlaid with the range data (using Garmin BaseCamp; Supplementary Table 2; Supplementary Figure 2). The locations of the transects in Waibira and Kaimira was based on a previous study (Hedges and McGrew, 2012; Supplementary Table 3-4).

We recorded the GPS start and end location, and we measured the width of the transect using a hip chain. To confirm that we were always following the straight transect line, we used a compass and the UTM location on the GPS.

### Processing faecal samples

Wearing rubber gloves, we half-filled each ziplock bag with water to soak the pieces, and we squashed big pieces to reduce the sample to a slurry. We then emptied the contents into a flat-bottomed sieve with a circular frame (diameter= 20cm, depth=8cm, mesh size= 1mm) over a large bucket. We sluiced the contents under running water, and then removed any extraneous foreign matter (such as twigs and dung beetles) with forceps. Following

Plumptre, Reynolds and Bakuneeta (1997), we counted the less abundant seeds in a sample, and estimated the number of very abundant seeds by counting the number in a sector of the sieve and then multiplying this number by the number of possible sectors in the sieve. We identified seeds with the aid of a “seed library” that we obtained by collecting the fruit of Known Fruit Trees and storing their seeds. If we were unable to identify an item at the time, I took a photo, described its characteristics, and cleaned it with ethanol to store it; all these seeds were eventually identified with the aid of experts. Some items were impossible to identify down to species, for example we recorded all fig species under *Ficus* spp. Once an item was identified and counted we removed it from the sieve. After all bags had been processed, we emptied remaining contents into the pit latrine to prevent contamination.

**Diversity analyses**

Diversity analysis in ecology is an incredibly complicated and controversial field, with an astonishing number of different indices, and debates over which one to use (Magurran, 2004). Whittaker (1960) divided the total species diversity at the regional or landscape scale (*gamma diversity γ*) into *alpha diversity* (*α*) and *beta diversity* (*β*). The former is the mean species diversity at the local, within-site scale, whereas the latter refers to the compositional turnover along a habitat gradient within one region (Cody, 1975; Whittaker, 1977). In more general terms, *α* diversity is the “effective number of species per compositional unit”, *β* diversity is the “number of compositional units in the dataset”, and *γ* diversity is the “total effective number of species in the dataset” (Tuomisto, 2010).

**Results – Known Fruit Trees analyses**

***i) Do the home ranges of the four communities differ in Known Fruit Tree abundance?***

The communities did not differ in abundance of Known Fruit Trees (GLMM, $\chi^2_{2,6} = 3.62$, $p = 0.16$).

#### *ii) Do Known Fruit Trees differ in size between the home ranges of the communities?*

The mean DBH of Known Fruit Trees differed between the communities' home ranges (GLMM, $\chi^2_{3,6} = 14.10$, $p < 0.005$).

For Known Fruit Trees, Sonso and Waibira had a higher mean DBH than Kaimira, and Sonso had a higher mean DBH than Mwera (Supplementary Table 6).

#### *iii) Does α, β and γ diversity of Known Fruit Trees differ between the communities' home ranges?*

The *α* diversity of Known Fruit Trees differed between the communities' home ranges (GLMM, $\chi^2_{2,6} = 10.60$, $p = 0.0051$), with Sonso having a diversity that was roughly twice as high as any other community (Supplementary Table 7). The *β* diversity of Known Fruit Trees did not differ between the communities' home ranges (GLMM, $\chi^2 2_{,6} = 5.64$, $p = 0.060$). The *γ* Known Fruit Tree diversity was roughly three times greater in Sonso than any other community (Supplementary Table 11).

#### *iv) Does the Known Fruit Tree species composition differ between the home ranges of the different communities?*

The Known Fruit Tree species composition differed between the community home ranges (ANOSIM, R= 0.25, p= 0.001).

The home ranges of Waibira and Kaimira had a similar species composition and relative abundance of Known Fruit Trees, whereas Sonso and Mwera differed in this respect (Supplementary Table 7).

**Supplementary Table 1.** The GPS coordinates of the transects in the Sonso community's home range, Budongo forest. There is trail system in the home range of the Sonso community and we thus noted which compartments were crossed during each transect.

| Transect number | GPS starting location | GPS finishing location | Compartments crossed |
|---|---|---|---|
| T1 | N1° 43.186' E31° 32.353' | N1° 43.185' E31° 32.084' | H6, I6, J6, L6, M6 |
| T2 | N1° 43.828' E31° 32.475' | N1° 43.828' E31° 32.205' | EF, FF, GF, HF, IF |
| T3 | N1° 44.182' E31° 32.828' | N1° 44.181' E31° 32.553' | (Extention line) |
| T4 | N1° 42.662' E31° 32.589' | N1° 42.661' E31° 32.320' | E16, G16, H16, I16, J16 |
| T5 | N1° 43.721' E31° 33.021' | N1° 43.721' E31° 32.751' | 5D, 4D, 3D, 2D, 1D |
| T6 | N1° 43.493' E31° 33.081' | N1° 43.493' E31° 32.811' | 60, 50, 40, 30, 20 |
| T7 | N1° 43.451' E31° 33.178' | N1° 43.180' E31° 33.178' | 80, 81, 81, 83, 84, 86 |
| T8 | N1° 42.989' E31° 32.335' | N1° 42.717' E31° 32.335' | H10, H11, H12, H13, H14, H15 |
| T9 | N1° 43.829' E31° 33.330' | N1° 43.558' E31° 33.330' | 11F, 11E, 11D, 11C, 11B |
| T10 | N1° 43.628' E31° 32.217' | N1° 43.356' E31° 32.217' | JB, JA, J0, J1, J2 |
| T11 | N1° 43.854' E31° 32.681' | N1° 43.583' E31° 32.681' | AG, AF, AE, AD, AC, AB |
| T12 | N1° 43.416' E31° 32.591' | N1° 43.145' E31° 32.591' | C2, C3, C4, C5a, C5b C6 |

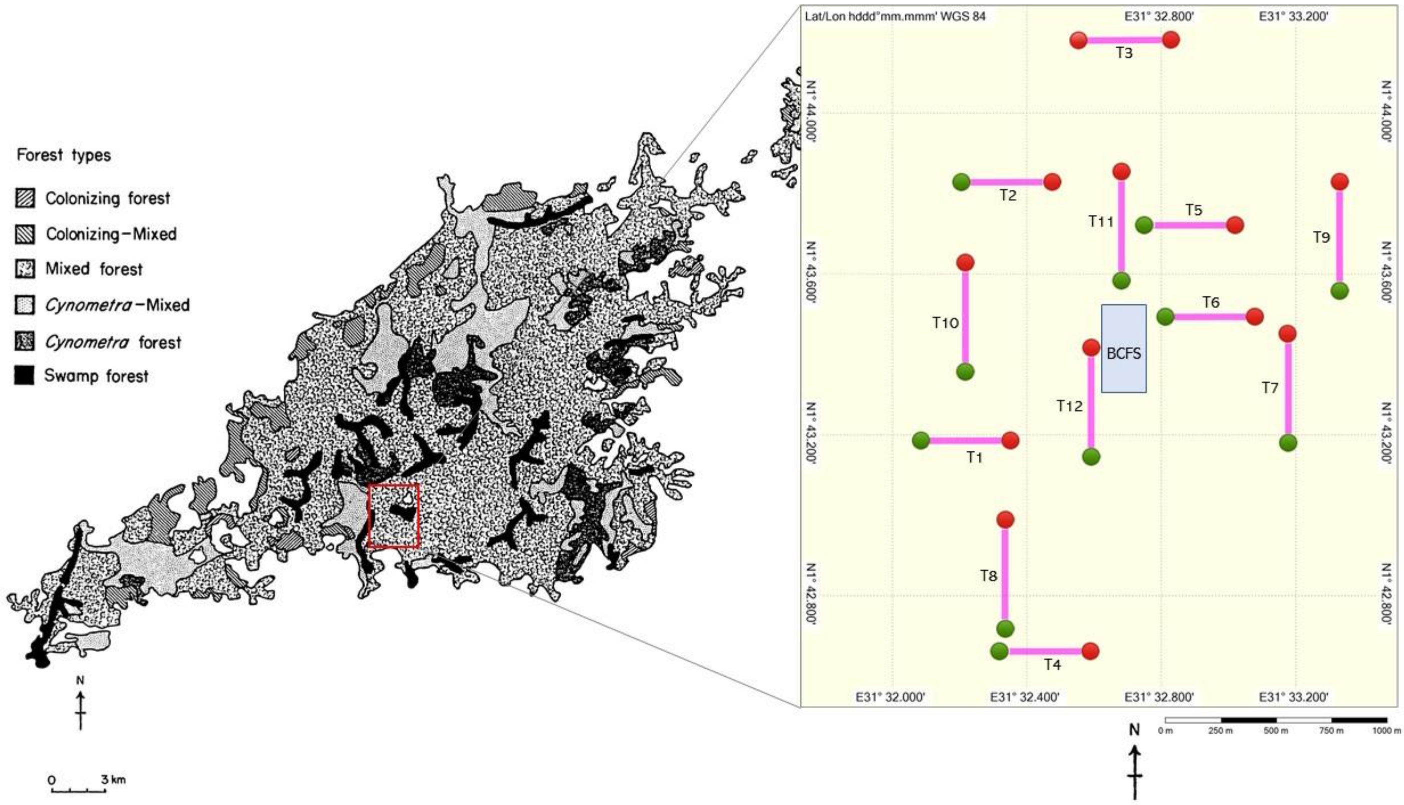


**Supplementary Figure 1.** Map of the transects in the Sonso community's home range (Budongo forest)

| Transect number | GPS starting location | GPS finishing location |
|---|---|---|
| T1 | N1° 18.713' E31° 00.482' | N1° 18.713' E31° 00.213' |
| T2 | N1° 19.355' E31° 00.603' | N1° 19.355' E31° 00.334' |
| T3 | N1° 19.709' E31° 00.957' | N1° 19.709' E31° 00.687' |
| T4 | N1° 18.189' E31° 00.718' | N1° 18.189' E31° 00.449' |
| T5 | N1° 19.248' E31° 01.150' | N1° 19.248' E31° 00.880' |

| | | |
|---|---|---|
| T6 | N1° 19.021' E31° 01.209' | N1° 19.020' E31° 00.939' |
| T7 | N1° 18.978' E31° 01.306' | N1° 18.707' E31° 01.306' |
| T8 | N1° 18.516' E31° 00.463' | N1° 18.245' E31° 00.464' |
| T9 | N1° 19.356' E31° 01.458' | N1° 19.085' E31° 01.459' |
| T10 | N1° 19.155' E31° 00.346' | N1° 18.884' E31° 00.346' |
| T11 | N1° 18.713' E31° 00.482' | N1° 18.713' E31° 00.213' |
| T12 | N1° 18.944' E31° 00.720' | N1° 18.672' E31° 00.720' |

**Supplementary Table 2.** The GPS coordinates of the transects in of the Mwera community's home range, Bugoma forest.

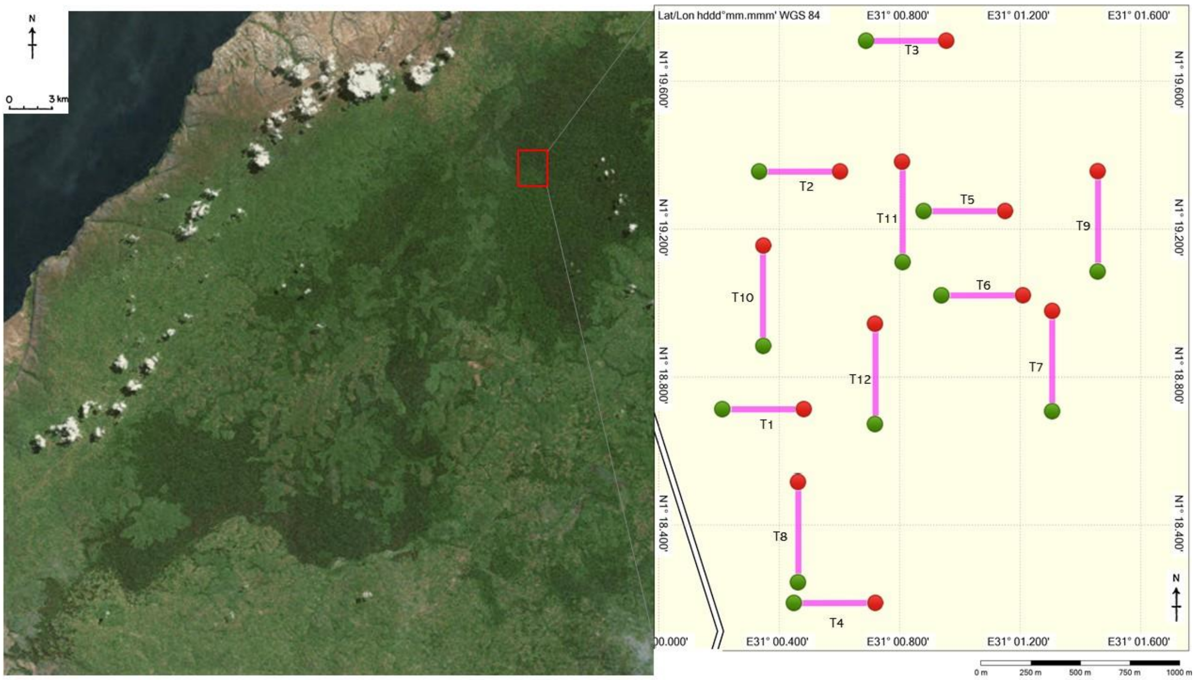


**Supplementary Figure 2**. Map of the transects in the Mwera community's home range (Bugoma forest).

**Supplementary Table 3.** The GPS coordinates of the transects in the home range of the Waibira community (Budongo Forest).

| Transect number | GPS starting location | GPS finishing location |
|---|---|---|
| 20F -> E | N1° 43.790’ E31° 33.803’ | N1° 43.773’ E31° 34.077’ |
| 27F -> N | N1° 43.763’ E31° 34.183’ | N1° 44.027’ E31° 34.185’ |
| 25D -> N | N1° 43.648’ E31° 34.075’ | N1° 43.923’ E31° 34.073’ |
| 30F -> E | N1° 43.752’ E31° 34.341’ | N1° 43.760’ E31° 34.609’ |
| 17L -> N | N1° 44.100’ E31° 33.645’ | N1° 44.367’ E31° 33.646’ |
| 25-2 -> N | N1° 43.322’ E31° 34.093’ | N1° 43.538’ E31° 34.084’ |
| 15R -> N | N1° 44.429’ E31° 33.526’ | N1° 44.700’ E31° 33.531’ |
| 15R -> E | N1° 44.429’ E31° 33.526’ | N1° 44.430’ E31° 33.799’ |
| 32N -> N | N1° 44.234’ E31° 34.451’ | N1° 44.498’ E31° 34.443’ |
| 25P -> E | N1° 44.362’ E31° 34.082’ | N1° 44.361’ E31° 34.351’ |

**Supplementary Table 4.** The GPS coordinates of the transects in the home range of the Kaimira community (Budongo Forest).

| Transect number | GPS starting location | GPS finishing location |
|---|---|---|
| JA -> W | N1° 43.537’ E31° 32.191’ | N1° 43.559’ E31° 31.922’ |
| JC -> W | N1° 43.624’ E31° 32.195’ | N1° 43.661’ E31° 31.926’ |
| JD -> W | N1° 43.704’ E31° 32.193’ | N1° 43.679’ E31° 31.922’ |
| JF -> W | N1° 43.828’ E31° 32.188’ | N1° 43.825’ E31° 31.918’ |
| JH -> W | N1° 43.945’ E31° 32.179’ | N1° 43.916’ E31° 31.906’ |
| JJ -> W | N1° 44.037’ E31° 32.187’ | N1° 44.007’ E31° 31.928’ |

| | | |
|---|---|---|
| OB -> N | N1° 43.568’ E31° 31.917’ | N1° 43.808’ E31° 31.956’ |
| QB -> N | N1° 43.591’ E31° 31.810’ | N1° 43.824’ E31° 31.818’ |
| RB -> N | N1° 43.590’ E31° 31.748’ | N1° 43.817’ E31° 31.771’ |
| TB -> N | N1° 43.585’ E31° 31.639’ | N1° 43.668’ E31° 31.649’ |

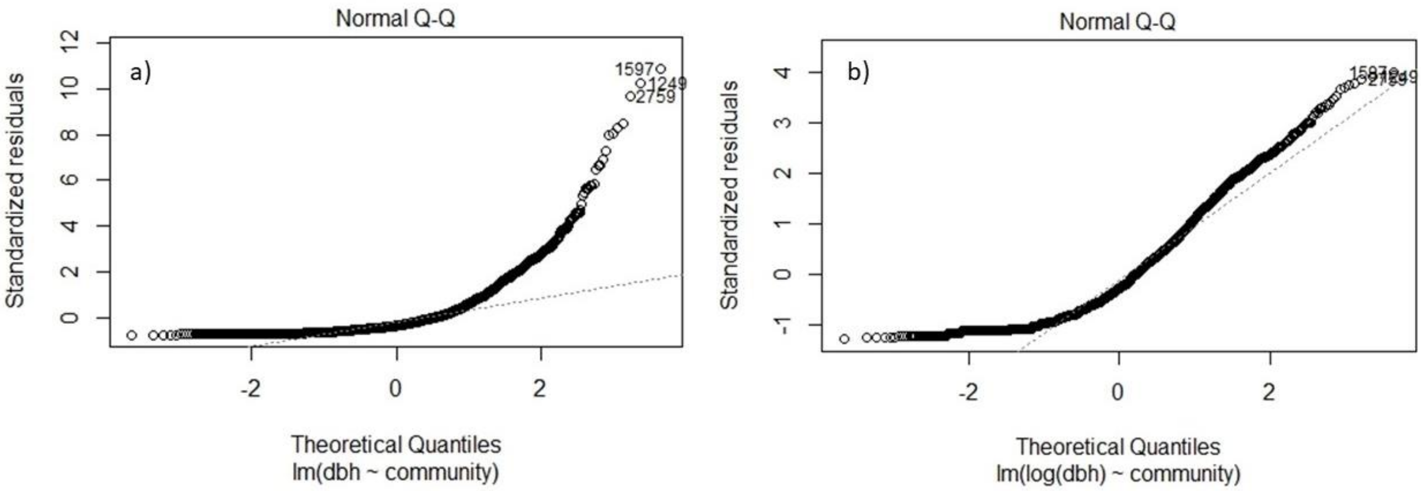


**Supplementary Figure 3**. Log-transformation on the diameter at breast height (DBH) measurements from the Known Feeding Trees studied in the home ranges of the different communities. The raw data of the DBH measurements did not meet the assumptions of normality (a), but after log-transformation the data was normally distributed (b) (albeit not perfectly).

**Supplementary Table 5.** Formulae of the diversity indices used to calculate alpha, beta and gamma diversity. Notations used: S = total number of species in the community; $p_i$ = proportion of S made up of the $i^{th}$ species; $x_{ij}$ = the number of individuals of species i in assemblage j; $x_{ik}$ = the number of individuals of species i in assemblage k; $N_j$ = total number of individuals in assemblage j; $N_k$ = total number of individuals in assemblage k.

| Diversity | Index | Formula | Hill number | Reference |
|---|---|---|---|---|
| **Alpha** **$H_\alpha$** | Shannon | $H' = -\sum_{i=1}^{S} p_i \ln p_i$ | $\exp(H_\alpha)$ | (Shannon, 1948) |
| **Beta** **$H_\beta$** | Horn | $\beta = 1 - \frac{\sum[(x_{ij} + x_{ik}) \log(x_{ij}x_{ik})] - \sum(x_{ij} \log x_{ij}) - \sum(x_{ik} \log}{[(N_j + N_k) \log(N_j N_k)] - (N_j \log N_j) - (N_k \log N_k}$ | $\exp(H_\beta)$ | (Horn, 1966) |
| **Gamma** **$H_\gamma$** | | $H_\gamma = H_\alpha + H_\beta$ | $\exp(H_\gamma) = \exp(H_\alpha) * \exp(H_\beta)$ | (Jost, 2006) |

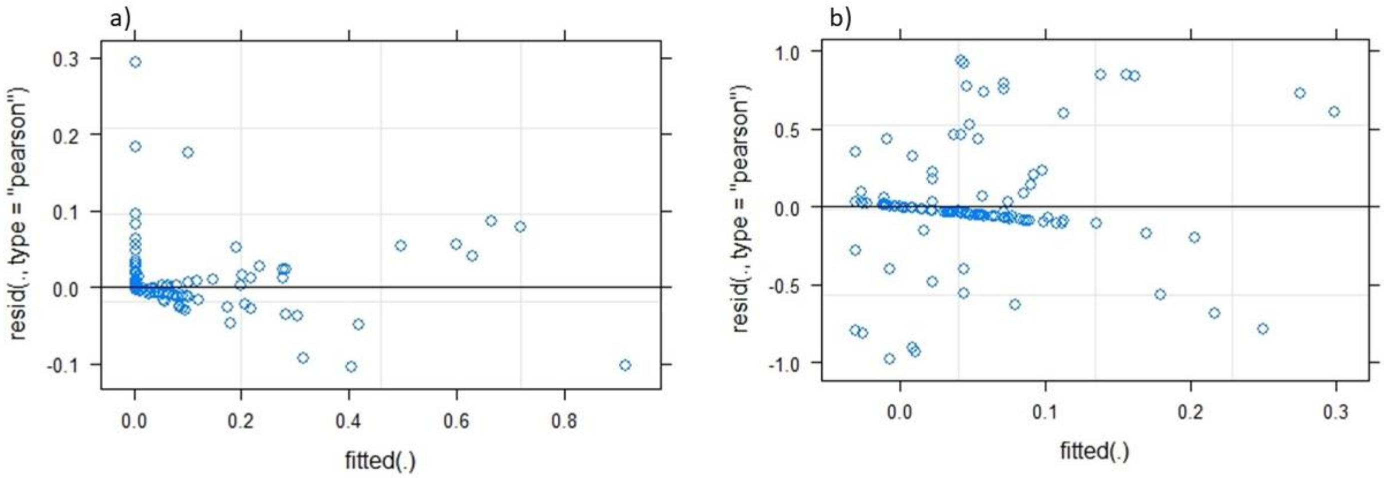


**Supplementary Figure 4.** Arcsin-squareroot transformation of the data on the proportion of seeds of a given plant species in the faecal samples. The raw data (a) did not meet the assumptions of homogeneity of variance and was thus arcsin-sqaureroot transformed (b) to meet the assumptions. This was done for the GLMM model looking at the predictability of faecal samples of chimpanzee diet.

**Supplementary Table 6.** Results of the Tukey test comparing the mean DBH of Known Feeding Trees and of Known Fruit Trees in the communities' home ranges.

| | Comparison | Difference | Lower CI | Upper CI | P-value |
|---|---|---|---|---|---|
| | Mwera-Kaimira | -1.25 | -3.43 | 0.92 | 0.42 |
| **Known Feeding Trees** | **Sonso-Kaimira** | **7.18** | **5.02** | **9.37** | **<0.001*** |
| | Waibira-Kaimira | -0.7 | -2.92 | 1.52 | 0.84 |
| | **Sonso-Mwera** | **8.45** | **6.32** | **10.58** | **<0.001*** |
| | Waibira-Mwera | 0.56 | -1.62 | 2.73 | 0.9 |
| | **Waibira-Sonso** | **-7.89** | **-10.07** | **-5.72** | **<0.001*** |
| | **Mwera-Kaimira** | **0.33** | **0.12** | **0.55** | **<0.001*** |
| **Known Fruit Trees** | **Sonso-Kaimira** | **0.31** | **0.09** | **0.52** | **<0.005*** |
| | **Waibira-Kaimira** | **0.25** | **0.028** | **0.48** | **0.021*** |
| | Sonso-Mwera | -0.026 | -0.23 | 0.18 | 0.99 |
| | Waibira-Mwera | -0.08 | -0.29 | 0.13 | 0.77 |
| | Waibira-Sonso | -0.054 | -0.27 | 0.16 | 0.92 |

**Supplementary Table 7.** Results of Tukey comparison of means of the alpha diversity of Known Fruit Trees and Known Feeding Trees between the communities' home ranges.

| | Comparison | Difference | Lower CI | Upper CI | P-value |
|---|---|---|---|---|---|
| **Known Feeding Trees** | Mwera-Kaimira | 0.088 | -0.0056 | 0.18 | 0.074 |
| | **Sonso-Kaimira** | **0.17** | **0.079** | **0.27** | **< 0.005 *** |
| | **Waibira-Kaimira** | **0.13** | **0.017** | **0.24** | **0.016 *** |
| | **Sonso-Mwera** | **0.086** | **0.0076** | **0.16** | **0.025 *** |
| | Waibira-Mwera | 0.041 | -0.058 | 0.14 | 0.71 |
| | Waibira-Sonso | -0.045 | -0.14 | -0.054 | 0.65 |
| **Known Fruit Trees** | Mwera-Kaimira | -0.04 | -0.12 | 0.042 | 0.59 |
| | **Sonso-Kaimira** | **0.081** | **0.0017** | **0.16** | **0.043 *** |
| | Waibira-Kaimira | -0.026 | -0.11 | 0.063 | 0.88 |
| | **Sonso-Mwera** | **0.12** | **0.051** | **0.19** | **< 0.005 *** |
| | Waibira-Mwera | 0.014 | -0.066 | 0.094 | 0.97 |
| | **Waibira-Sonso** | **-0.11** | **-0.18** | **-0.029** | **< 0.005 *** |

**Supplementary Table 8.** Indicator values (in percentage) for Known Feeding Tree species at the home ranges of the different communities.

| Communities | Known Feeding Tree species | IndVal (%) | P-value |
|---|---|---|---|
| Sonso | *Trichilia rubescens* | 99.1 | 0.001 |
| | *Teclea nobilis* | 91.3 | 0.001 |
| | *Croton sylvaticus* | 87.9 | 0.001 |
| | *Entandrophragama angolense* | 86.6 | 0.001 |
| | *Milicia excelsa* | 66.1 | 0.003 |
| | *Erythrophleum suaveolens* | 64.5 | 0.002 |
| | *Cleistopholis patens* | 57.7 | 0.014 |
| | *Macaranga schweinfurthii* | 57.7 | 0.010 |
| | *Mammea africana* | 53.4 | 0.017 |

| | | | |
|---|---|---|---|
| | *Trichilia dregeana* | 50.0 | 0.050 |
| Mwera | *Morus lactea* | 79.5 | 0.001 |
| | *Chrysophyllum muerense* | 76.4 | 0.001 |
| | *Sterculia dawei* | 50.0 | 0.046 |
| Sonso + Kaimira | *Khaya anthoteca* | 93.5 | 0.001 |
| Sonso + Mwera | *Myrianthus holstii* | 86.6 | 0.001 |
| | *Celtis wightii* | 84.2 | 0.001 |
| | *Caloncoba schweinfurthii* | 76.6 | 0.010 |
| | *Antiaris toxicaria* | 75.6 | 0.001 |
| | *Ficus variifolia* | 73.6 | 0.001 |
| | *Ricinodendron heudelotii* | 64.5 | 0.006 |
| | *Desplatsia dewevrei* | 61.2 | 0.011 |
| Sonso + Waibira | *Ficus exasperata* | 70.8 | 0.006 |
| | *Cola gigantea* | 55.2 | 0.035 |

**Supplementary Table 9.** Indicator values (in percentage) for Known Fruit Tree species at the home ranges of the different communities.

| Communities | Known Feeding Tree species | IndVal (%) | P-value |
|---|---|---|---|
| Sonso | *Croton sylvaticus* | 87.9 | 0.001 |
| | *Milicia excelsa* | 66.1 | 0.001 |
| | *Cleistopholis patens* | 57.7 | 0.005 |
| | *Macaranga schweinfurthii* | 57.7 | 0.010 |
| | *Mammea africana* | 53.4 | 0.024 |
| | *Strychnos mitis* | 50.0 | 0.046 |
| Mwera | *Morus lactea* | 79.5 | 0.001 |
| | *Chrysophyllum muerense* | 76.4 | 0.001 |
| Sonso + Mwera | *Myrianthus holstii* | 86.6 | 0.001 |
| | *Caloncoba schweinfurthii* | 76.6 | 0.012 |
| | *Antiaris toxicaria* | 75.6 | 0.001 |
| | *Ficus variifolia* | 73.6 | 0.001 |
| | *Desplatsia dewevrei* | 61.2 | 0.006 |
| Sonso + Waibira | *Ficus exasperata* | 70.8 | 0.005 |
| | *Cola gigantea* | 55.2 | 0.033 |

**Supplementary Table 10.** Results of the Tukey test comparing the mean of the abundance of insect nests between the home ranges of the four communities.

| Comparison | Difference | Lower CI | Upper CI | P-value |
|---|---|---|---|---|
| **Mwera-Kaimira** | **-8.72** | **-16.1** | **-1.34** | **0.015** |
| **Sonso-Kaimira** | **-13.47** | **-20.85** | **-6.09** | **<0.001** |
| Waibira-Kaimira | -5.65 | -13.18 | 1.9 | 0.2 |
| Sonso-Mwera | -4.75 | -11.97 | 2.47 | 0.31 |
| Waibira-Mwera | 3.08 | -4.3 | 10.46 | 0.68 |
| **Waibira-Sonso** | **7.83** | **0.45** | **15.21** | **0.034** |

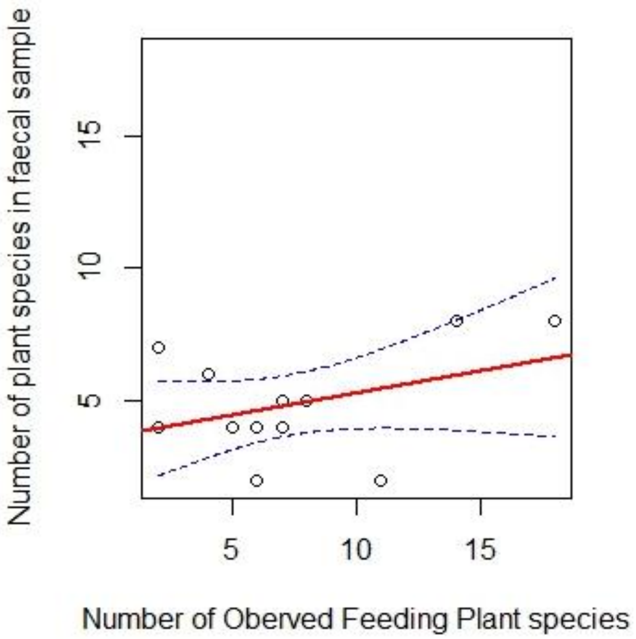


**Supplementary Figure 5.** The number of plant species recorded in the faecal samples increases with increasing number of Observed Feeding Plant species. The slope and the 95% confidence intervals were estimated from the model fitted.

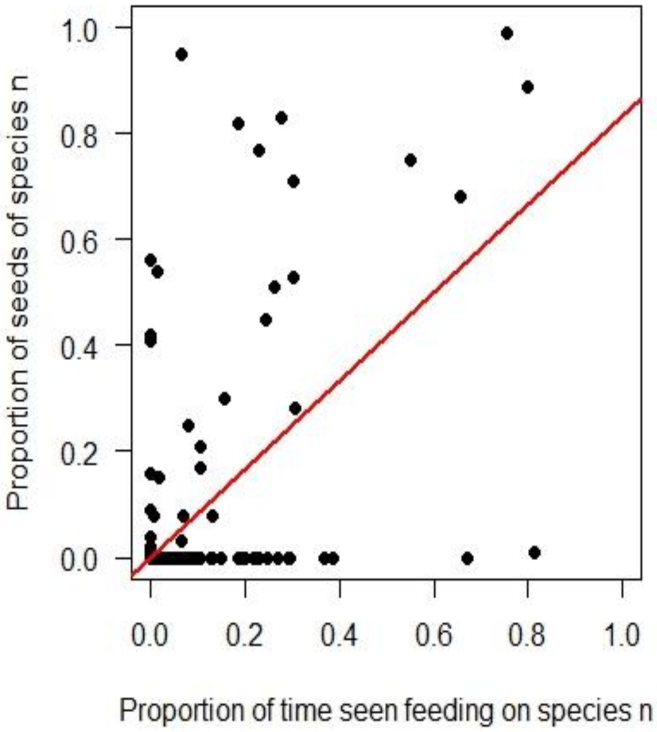


**Supplementary Figure 6.** The proportion of seeds of a particular plant species increases with the proportion of time we observed the Sonso chimpanzees to feed on that species. The slopes were predicted from the respective GLMM on the arcsine square root transformed data, with the coefficient then back-transformed to the original data.

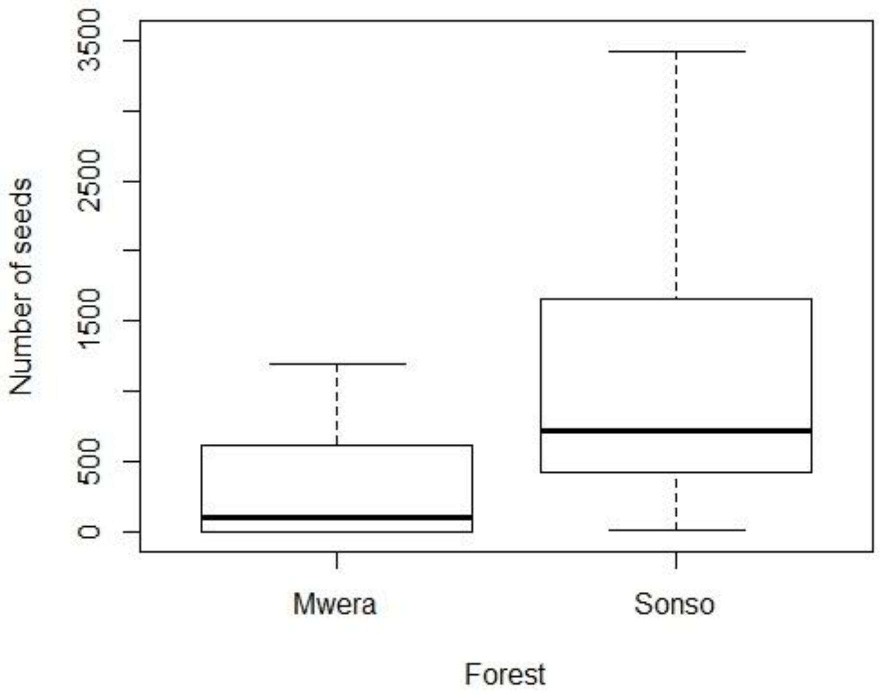


**Supplementary Figure 7.** Boxplot illustrating the median and the upper and lower quartiles of the abundance of seeds per faecal sample for each chimpanzee community.

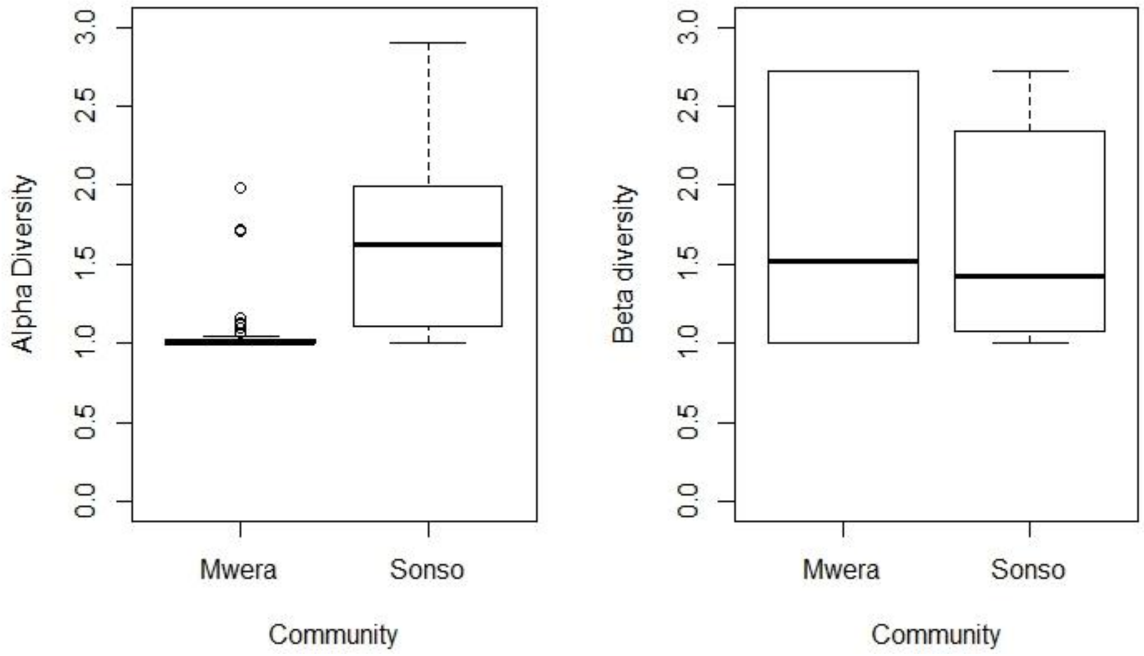


**Supplementary Figure 8.** Boxplot illustrating the median and the upper and lower quartiles of (a) alpha and (b) beta diversities of the seeds in the faecal samples of the Mwera and Sonso communities.

**Supplementary Table 11.** Diversity indices and Alpha, Beta and Gamma diversity of Known Fruit Tree[illegible] (a) ach community's range. The communitie[illegible] (b) the Budongo forest are shaded grey.

| | Known Fruit Trees | | | | |
|---|---|---|---|---|---|
| | **Shannon-Wiener** | **Horn** | **α** | **β** | **Γ** |
| Sonso | 2.03 | 0.43 | 7.89 | 1.60 | 15.38 |
| Mwera | 1.24 | 0.34 | 3.67 | 1.56 | 5.71 |
| Waibira | 1.37 | 0.30 | 4.05 | 1.36 | 5.51 |
| Kaimira | 1.19 | 0.13 | 3.41 | 1.15 | 3.92 |
